# Modeling the short-term dynamics of *in vivo* excitatory spike transmission

**DOI:** 10.1101/475178

**Authors:** Abed Ghanbari, Naixin Ren, Christian Keine, Carl Stoelzel, Bernhard Englitz, Harvey A. Swadlow, Ian H. Stevenson

## Abstract

Information transmission in neural networks is influenced by both short-term synaptic plasticity (STP) as well as non-synaptic factors, such as after-hyperpolarization currents and changes in excitability. Although these effects have been widely characterized *in vitro* using intracellular recordings, how they interact *in vivo* is unclear. Here we develop a statistical model of the short-term dynamics of spike transmission that aims to disentangle the contributions of synaptic and non-synaptic effects based only on observed pre- and postsynaptic spiking. The model includes a dynamic functional connection with short-term plasticity as well as effects due to the recent history of postsynaptic spiking and slow changes in postsynaptic excitability. Using paired spike recordings, we find that the model accurately describes the short-term dynamics of *in vivo* spike transmission at a diverse set of identified and putative excitatory synapses, including a thalamothalamic connection in mouse, a thalamocortical connection in a female rabbit, and an auditory brainstem synapse in a female gerbil. We illustrate the utility of this modeling approach by showing how the spike transmission patterns captured by the model may be sufficient to account for stimulus-dependent differences in spike transmission in the auditory brainstem (endbulb of Held). Finally, we apply this model to large-scale multi-electrode recordings to illustrate how such an approach has the potential to reveal cell-type specific differences in spike transmission *in vivo*. Although short-term synaptic plasticity parameters estimated from ongoing pre- and postsynaptic spiking are highly uncertain, our results are partially consistent with previous intracellular observations in these synapses.

**Significance Statement:** Although synaptic dynamics have been extensively studied and modeled using intracellular recordings of post-synaptic currents and potentials, inferring synaptic effects from extracellular spiking is challenging. Whether or not a synaptic current contributes to postsynaptic spiking depends not only on the amplitude of the current, but also on many other factors, including the activity of other, typically unobserved, synapses, the overall excitability of the postsynaptic neuron, and how recently the postsynaptic neuron has spiked. Here we developed a model that, using only observations of pre- and postsynaptic spiking, aims to describe the dynamics of *in vivo* spike transmission by modeling both short-term synaptic plasticity and non-synaptic effects. This approach may provide a novel description of fast, structured changes in spike transmission.

## Introduction

In response to a presynaptic input, the amplitudes of elicited postsynaptic potentials (PSPs) can increase or decrease dramatically due to short-term synaptic plasticity (Zucker and Regehr, 2002; Regehr, 2012). The probability that a postsynaptic neuron spikes in response to a presynaptic spike can also increase or decrease depending on the recent history of pre- and postsynaptic activity (Usrey et al., 2000; Swadlow and Gusev, 2001). Although many models exist to describe intracellular observations of short-term synaptic plasticity (Costa et al., 2013; Hennig, 2013; Barri et al., 2016; Bird et al., 2016), most models of *functional* connections between neurons based on extracellular spike observations assume that connections are fixed over time (Truccolo et al., 2005; Pillow et al., 2008). Unlike intracellular PSP observations, where the amplitude of each individual presynaptic spike can be measured (subject to noise), extracellular spike observations are sparse, typically all-or-none binary events. Modeling dynamic, functional connections from spike observations, especially in the presence of uncontrolled, ongoing neural activity, presents a major statistical challenge (Ghanbari et al., 2017). Here we further develop a model-based approach that, given only pre- and postsynaptic spike observations, estimates the contributions of short-term synaptic plasticity and several non-synaptic factors to the probability of spike transmission.

Traditionally, the influence of presynaptic spikes on postsynaptic spiking is measured using cross-correlation (Perkel et al., 1967; Fetz et al., 1991; Csicsvari et al., 1998; Barthó et al., 2004). If two neurons are monosynaptically connected, the probability of the postsynaptic neuron spiking will briefly increase or decrease following a presynaptic spike, which appears as a fast-onset, short-latency peak or trough in the cross-correlation, depending on whether the synapse is excitatory or inhibitory (Perkel et al., 1967; Barthó et al., 2004). Just as synaptic potentials depress or facilitate due to short-term synaptic plasticity, this spike transmission probability might also depend on the recent history of presynaptic activity. By subdividing cross-correlograms to characterize the specific effects of different presynaptic spike patterns, previous studies have found that certain, putative synaptic connections show reduced spike transmission probability following recent presynaptic spikes (Swadlow and Gusev, 2001; English et al., 2017), while others show increased probability (Usrey et al., 2000), as might be expected of depressing or facilitating synapses, respectively.

Here, rather than subdividing correlograms, we use a likelihood-based modeling approach that extends previous static models of functional connectivity (Harris et al., 2003; Pillow et al., 2008; Stevenson et al., 2008). This dynamic model describes not only the sign and strength of synaptic connections, but also whether the dynamics are depressing or facilitating. In addition to describing differences in responses to specific presynaptic spike patterns, the model-based approach also allows us to predict how the postsynaptic neuron will respond to arbitrary patterns of presynaptic activity. In previous work, we evaluated this type of dynamical functional connectivity model on simulated and *in vitro* experiments where the ground-truth dynamics were known (Ghanbari et al., 2017). These results demonstrated that, at least in a controlled setting, short-term synaptic plasticity can be inferred from spike observations, even in the presence of sources of error, such as spike sorting errors, stochastic vesicle release, and common input from unobserved neurons. Here we build on this model and examine how well it can account for excitatory spike transmission dynamics observed *in vivo* where the true synaptic currents are unknown.

A key element of our dynamical functional connectivity model is the inclusion of both synaptic and non-synaptic contributions to spike transmission. For each individual presynaptic spike, our model predicts postsynaptic spiking by taking into account synaptic coupling with STP, synaptic summation, post-spike history effects, and slow fluctuations of excitability. Although these effects do not include all factors that may influence spiking statistics (Herz et al., 2006), together they can account for wide variety of phenomena, including subthreshold membrane integration (Carandini et al., 2007) and slower fluctuations in the overall excitability of the postsynaptic neuron, such as observed during neuromodulation (Henze and Buzsáki, 2001). The interaction between synaptic and non-synaptic effects, as well as the degree to which each factor contributes is likely to lead to diverse patterns of spike transmission. Here we show how models of dynamical functional connectivity with short-term synaptic plasticity can capture these patterns of spike transmission and disentangle the multiple factors that shape postsynaptic response.

## Material and methods

### Neural Data

All data analyzed here were obtained from previous studies (see below). Animal use procedures were approved by the institutional animal care and use committees at University of Connecticut (VB-Barrel), University of Leipzig (ANF-SBC), or University College London (MEA), respectively, and conform to the principles outlined in the Guide for the Care and Use of Laboratory Animals (National Institutes of Health publication no. 86-23, revised 1985).

To illustrate how synaptic dynamics can be estimated from spikes, we first examined a set of three strong putative or identified synapses with diverse spike transmission probability patterns: (i) a local, excitatory connection from one neuron in mouse thalamus to another detected from a larger multi-electrode array (MEA) recording, (ii) a ventrobasal thalamus projection to primary somatosensory cortex (VB – Barrel) in a rabbit, and (iii) an *in vivo* loose-patch (juxtacellular) recording of an auditory nerve projection onto a spherical bushy cell (ANF-SBC) in the auditory brainstem of a gerbil. We then use this auditory brainstem connection to explore how synaptic transmission probability depends on the stimulus and compare the results with a model without short-term synaptic plasticity. Next, we applied our model more generally to analyze a large sample of putative synaptic connections recorded from the MEA dataset. The data from these three identified strong synapses and the MEA data have been collected from different species, regions, cell-types, under different stimulation and show a diverse pattern of postsynaptic spiking probability. In all cases we deduce short-term synaptic dynamics on the basis of only pre- and postsynaptic spike observations.

For the first putative synapse, we use *in vivo* data from simultaneous extracellular recordings in ventrobasal (VB) thalamic barreloids and topographically aligned, somatosensory cortical barrel columns (VB-Barrel) in awake, unanesthetized, adult rabbits. Detailed surgical and physiological methods have been described previously (Swadlow and Gusev, 2002). Spike-triggered averages of the cortical spikes following spiking of the VB neuron was used to identify connected S1 neurons. Based on the presence of high frequency discharge (3+ spikes, > 600 Hz) following electrical stimulation of the thalamus, and narrow spike waveforms, the S1 neuron in this recording was identified as a putative inhibitory neuron (Kawaguchi, 2001). These recordings identified several putative thalamocortical projections. The putative synapse that we model here is particularly clear, with 68,345 pre- and 128,096 postsynaptic spikes recorded over the course of 92 minutes of spontaneous activity and has been previously studied in (Swadlow and Gusev, 2001; Swadlow, 2002).

For the second synapse, we examined *in vivo* loose-patch recordings at the Endbulb of Held in young adult female gerbils. Detailed surgical and physiological methods have been previously described (Keine et al., 2017). Briefly, the glass electrode was positioned in the anterior portion of the ventral cochlear nucleus (AVCN) and single-units were recorded during varying acoustic stimulation. Single units were classified when recording a positive action potential amplitude of at least 2 mV and showing the characteristic complex waveform identifying them as large spherical bushy cells (SBC) of the rostral AVCN. This recording included a mixture of juxtacellular waveforms: an isolated excitatory PSP (EPSP) or an EPSP followed by a postsynaptic action potential. For both cases the timing of EPSPs and spikes and rising slope of the EPSPs were extracted. The timing and slope of the EPSPs were identified using a slope threshold for the rising part of EPSPs as previously described (Keine et al., 2016). We then modeled spike transmission probability patterns for two recordings: (i) during randomized pure tone acoustic stimulation and (ii) during multiple stimuli, i.e. randomized frequency-level pure tone stimulation interspaced with spontaneous activity, natural sounds, and also during spontaneous activity. Using this second dataset, we characterized how variable presynaptic spike patterns evoked by different stimuli affected the patterns of spike transmission at the same synapse.

We also use MEA spiking data to study the factors shaping spike transmission probability patterns in a large-scale recording with multiple cell-types. Here we use a previously collected, publicly available recording from the Cortex Lab at UCL (Jun et al., 2017; Mora Lopez et al., 2017) with data from two Neuropixels electrode arrays recorded simultaneously, each with 960 sites (384 active) with lengths of 10-mm and spacing of 70 × 20-μm (http://data.cortexlab.net/dualPhase3/). The two electrode arrays span multiple brain areas and ∼90 min of data was collected in an awake, head-fixed mouse on a rotating rubber wheel during visual stimulus presentations. Spikes were automatically detected and sorted using Kilosort (Pachitariu et al., 2016) on the broadband (0.3– 10 kHz) signal and then manually curated. If two clusters of spikes had similar waveforms, cross-correlogram features, and spike amplitudes, they were merged into a single cluster and assigned to a single neuron. In total, 831 well-isolated single neurons where identified from the two probes in several different brain areas: visual cortex (n=74), hippocampus (n=64), thalamus (n=244), motor cortex (n=243), and striatum (n=200). Due to the large number of simultaneously recorded neurons in this dataset, there are many potential synapses (∼831^2^).

### Synapse Detection

To identify putative monosynaptic connections between well-isolated single neurons, we looked for specific patterns in the cross-correlograms (Moore et al., 1970). If two neurons are monosynaptically connected, the probability of postsynaptic spiking increases/decreases rapidly following a presynaptic spike. In spiking data, this rapid, transient change can be seen in cross-correlograms as an asymmetric bump/dip in the number of postsynaptic spikes following presynaptic spikes (Barthó et al., 2004). For each connection we calculated the cross-correlogram in a 5 ms window before and after presynaptic spikes with bin-size of 0.1 ms. To avoid aliasing in the cross-correlograms, we added a small, random shift to each postsynaptic spike drawn uniformly between −Δt/2 and Δt/2 where Δt is the spike time resolution (0.01 ms in most cases). Here we used a model-based approach using the cross-correlograms to decide whether two synapses are monosynaptically connected. To fit the cross-correlogram we used a baseline rate *μ*, a linear combination of B-spline bases **B**(t), and a weighted alpha function to model the synapse, *w α*(t), all passed through an output nonlinearity; *λ*(*t*) = exp(*μ* + ***r*****B**(t) + *w α*(t)). The alpha function, *α*(t) = (t – *t*_*d*_)/*τ*_*α*_ exp(1 − (t − *t*_*d*_)/*τ*_*α*_), describes the shape of the synaptic potential where *t*_*d*_ is the synaptic delay and *τ*_*α*_ is the synaptic time-constant (Carandini et al., 2007). For individual connections, we estimate these parameters by maximizing the penalized Poisson log-likelihood *l*(*μ, **r**, w, t*_*d*_, *τ*_*α*_) = Σy_i_*logλ*_*i*_ − Σ*λ*_*i*_ + ϵ‖*r*‖_2_ where *y*_*i*_ is the number of postsynaptic spikes observed in the *i*-th bin of the correlogram and ‖ *r*‖_2_ regularizes the model to penalize B-spline bases for capturing sharp increases in the cross-correlogram. *ϵ* is a regularization hyper-parameter which we set to 1 based on manual search. Due to the parameterization of *α*(t), the log-likelihood is not concave. However, since the gradient of the log-likelihood can be calculated analytically, we efficiently optimize the likelihood using a gradient-based pseudo-Newton method (LBFGS) (Boyd and Vandenberghe, 2004). During the optimization, the delay and time-constant parameters are log-transformed, allowing us to use unconstrained optimization, even though they are strictly positive. We used random restarts to avoid local maxima. To identify putative monosynaptic connections in the large-scale multi-electrode array data, we compared this model with a smooth model with slow changes in cross-correlogram and without the synapse, *λ*_0_(t) = exp(*μ*′ + ***r*****′B**(t)), using the log-likelihood ratio (LLR) test between our full model with synapse and the nested smooth model. Since low values of the likelihood ratio mean that the observed result was better explained with full model as compared to the smooth model, we then visually screened pair-wise connections with lowest ratios (LLR <-6) compared to the null model to find putative synapses. Out of ∼831^2^ possible connections in this dataset we find ∼200 putative synapses (0.0*3*%). We handpicked a strong putative synapse between two thalamic neurons to study its efficacy pattern in detail alongside the VB-Barrel and ANF-SBC synapses.

In addition to this single strong synapse, we also categorize putative pre- and postsynaptic cell types for the connections detected in the MEA dataset. For this purpose, we assessed single units based on their cross-correlograms, firing rates, and spike waveforms. We categorized units as excitatory or inhibitory if, in accordance with Dale’s law, all outgoing cross-correlograms showed transient, short-latency (<4ms) increase/decrease in spiking probability. We then looked into identified inhibitory neurons and categorized them into to putative fast-spiking (FS) and regular-spiking (RS) inhibitory neurons. Using these putative Excitatory-FS and Excitatory-RS synapses, we then examine how the spike transmission patterns differ for these two subtypes of inhibitory neurons.

### Extending a Generalized Linear Model to Account for Short-term Plasticity (TM-GLM)

Short-term synaptic plasticity causes the amplitude of postsynaptic potentials (PSP) to vary over time depending on the dynamics of synaptic resources and utilization and can be modeled using the pattern of presynaptic spiking (Markram et al., 1998; Tsodyks et al., 1998). However, changes in the overall postsynaptic spiking probability cannot be uniquely attributed to changes in amplitudes of postsynaptic potentials. To accurately describe the dynamics of spike transmission, we also need to account for the membrane potential summation, the excitability of the postsynaptic neuron (e.g. slow changes in the presynaptic firing rate) and the dynamics of postsynaptic spiking (e.g. refractory period, after hyperpolarization current). We developed an extension of a generalized linear model, which we call a TM-GLM to describe each of these effects. Concretely, the probability of a postsynaptic spike shortly after each presynaptic spike accounts for the full sequence of previous presynaptic spiking and the recent history of postsynaptic spiking. We define the conditional intensity of the postsynaptic neuron after the *i*-th presynaptic spike, 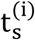, so that the probability of observing a postsynaptic spike in the *j*-th time bin after the *i*-th presynaptic spike is given as:

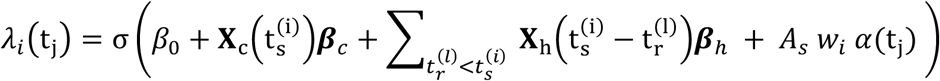

where 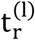 are the postsynaptic spike times preceding 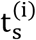. For each presynaptic spike, our model decomposes the firing rate of the postsynaptic neuron into four effects: a baseline firing rate, *β*_0_, slow fluctuations in postsynaptic firing rate **X**_**c**_***β***_***c***_, history effects from the recent postsynaptic spikes (prior to 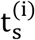), **X**_**h**_***β***_***h***_, and a time-varying coupling effect from the presynaptic input, *A*_*s*_*w α*(*t*) (Fig. 1).

**Fig. 1:**
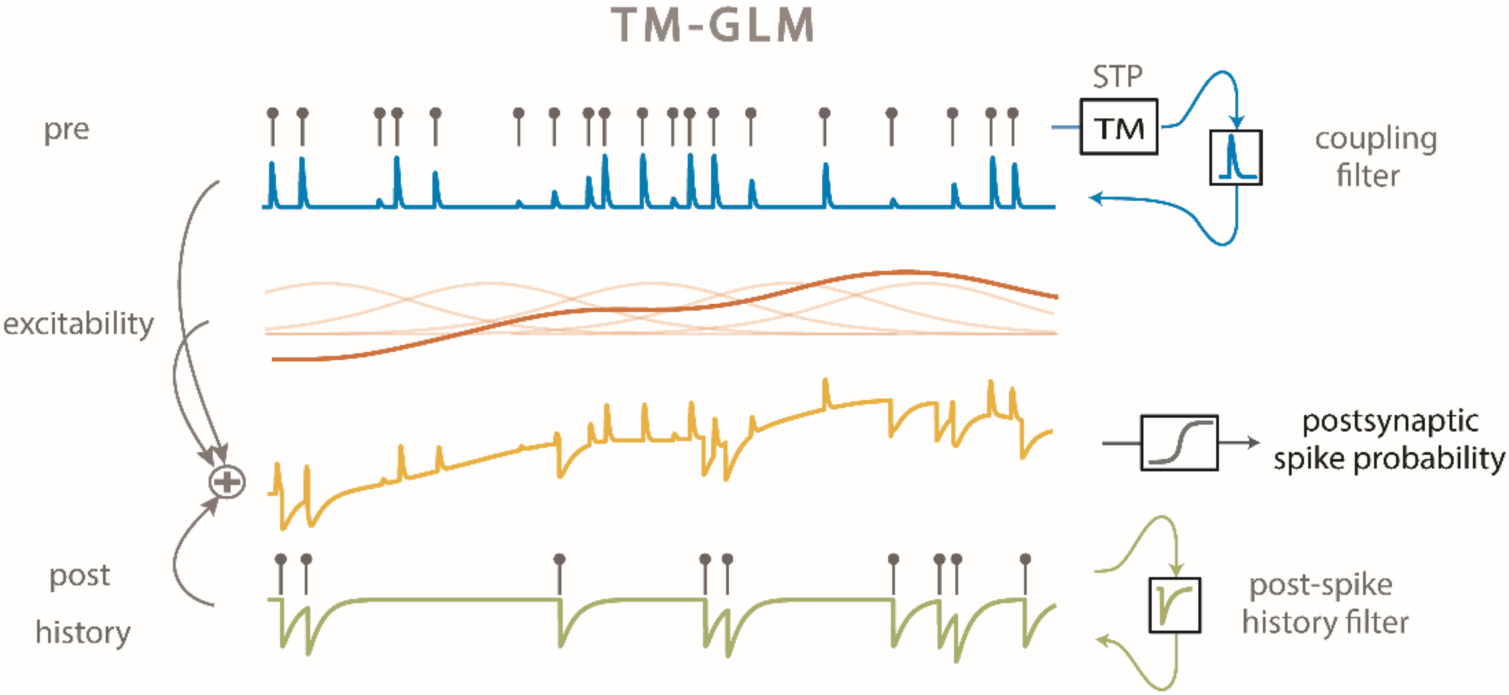
TM-GLM. Postsynaptic spiking probability before passing the spiking nonlinearity (yellow) changes as a linear combination of presynaptic coupling term with STP dynamics (blue), postsynaptic spiking history (green), the postsynaptic excitability (red). Transparent red curves show the bases of slow changes in postsynaptic probability at presynaptic spike times (*X*_*c*_).

Here we model slow fluctuations in the postsynaptic rate **X**_**c**_*β*_*c*_ with a linear combination of B-splines with equally spaced knots every 50 seconds of recording time. In the history term, splines (**X**_**h**_) span a period of 10 ms prior to each presynaptic spike with 4 logarithmically-spaced knots. By scaling *α*(t_j_) with a multiplicative factor, *w*_*i*_, the strength of a synapse can vary over time and, in this case, depends on the detailed sequence of presynaptic spiking and their corresponding inter-spike intervals. *A*_*s*_ is the magnitude of the synaptic strength. In this case we use a model for short-term synaptic plasticity that allows both depression (where the *w*_*i*_ decreases for shorter presynaptic ISIs) and facilitation (where the *w*_*i*_ increases for shorter presynaptic ISIs), and incorporates membrane summation. To model these effects, *w*_*i*_ is determined by a nonlinear dynamical system based on the Tsodyks and Markram (TM) model (Tsodyks and Markram, 1997; Markram et al., 1998) where: 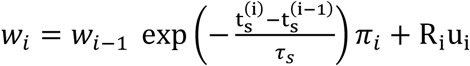, where *τ*_*s*_ is the membrane time-constant and the first term of the equation describes how postsynaptic membrane potential summation increases the probability of postsynaptic spiking. This membrane summation will be ignored if there is a postsynaptic spike: 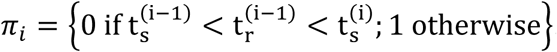. In the second term of this equation, *R* represents the dynamics of resources and *u* describes their utilization.

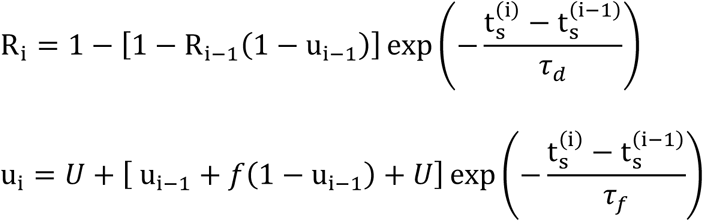

where *τ*_*d*_ and *τ*_*f*_ are the depression and facilitation time-constants. *U* is the release probability, and *f* is the magnitude of facilitation. To make the estimation more tractable, we approximate the full optimization problem and estimate synaptic delay, *t*_*d*_, and time-constant, *τ*_*α*_, by fitting *α*(*t*) using the full cross-correlogram, as above. We fix these parameters for the rest of the optimization process. We then maximize a penalized, Bernoulli log-likelihood log (*l*(*θ*)) = ΣΣ [y_ij_*λ*_*i*_ (t_j_)− (1 − y_ij_) (1 − *λ*_*i*_(t_j_))] + *γ*‖**θ′** _*stp*_‖_2_ where *γ* = 1 is the regularization hyperparameter to estimate the parameters: ***θ*** = {*β*_0_, ***β***_*c*=1;*C*_, ***β***_*h*=1:*H*_, *A*_*s*_, ***θ***_*stp*_}, ***θ***_*stp*_ = {*τ*_*d*_, *τ*_*f*_, *U, f, τ*_*s*_}.

As with previous applications of GLMs, we assume that bins are conditionally independent given the covariates, but unlike many other GLMs, here we only calculate the log-likelihood during short intervals (5ms) after presynaptic spikes. With y_ij_ being a binary value representing the presence of a postsynaptic spike in the *j*-th time bin after the *i*-th presynaptic spike. We again used a logarithmic transformation for the time-constants to avoid negative values and logit transformation for *U* and *f* to bound their values in the interval [0, 1]; ***θ***′_*stp*=_ = {log (*τ*_*d*_), log (*τ*_*f*_), log it(*U*), log it(*f*), log (*τ*_*s*_)}. By modeling STP this model is no longer a strict GLM, and the log-likelihood may have local maxima. Here we use random restarts to avoid local maxima in our optimization process. The parameters of each restart {*β*_0_, ***β***_*c*=1;*C*_, ***β***_*h*=1:*H*_, *A*_*s*_} are initialized by adding noise (∼ *N*(0,1)) to the corresponding parameters in a standard GLM. We initialize the log-transformed plasticity parameters with 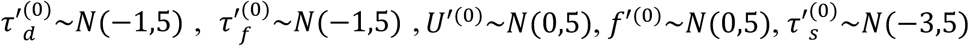. We then use an LBFGS algorithm to optimize the log-likelihood where we calculate all derivatives analytically except for derivatives of *θ*_*stp*_ which we calculate numerically. To estimate the uncertainty of the parameters, we bootstrap the data from each of the strong synapses by chunking the whole recording time into samples of 50 seconds then resampling the chunks to generate a new spike train with the same original length.

### Calculating spike transmission probability

To demonstrate how the probability of postsynaptic spiking changes according to the corresponding presynaptic inter-spike intervals, we estimated spike transmission probabilities from the cross-correlograms directly instead of using a model. To calculate this probability, we focused on a transmission interval after the presynaptic spike where the conditional intensity (when corrected for the baseline rate) goes above 10% of the maximum of ***α***(*t*) (horizontal bars in Fig 2A). We split the presynaptic inter-spike interval distribution into log-spaced intervals, and, for each interval, we calculate the ratio between numbers of postsynaptic spikes in the transmission interval to the number of presynaptic spikes. Unlike previous studies (Swadlow and Gusev, 2001, 2002) we do not correct this probability for the baseline postsynaptic rate. The uncorrected probability allows us to more directly compare the model predictions to the empirical spike transmission probabilities. Since our model gives an estimate of the postsynaptic probability after each individual presynaptic spike, we can average over the same transmission interval. However, we know if there is a postsynaptic spike in the transmission interval, probability of a postsynaptic spike goes to ∼0 for all consecutive bins due to the post-spike dynamics (e.g. refractory period). Therefore, we measure the predicted probability of a postsynaptic spike in a 5ms window after *i*-th presynaptic spike from binned *λ*_*i*_(t_j_) as follows: 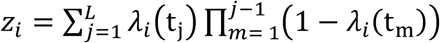. Here we assume conditional independence of the *j*-th bin after a presynaptic spike, but we enforce a refractory period for all bins after a postsynaptic spike in our generative model. Here *L* is the first bin that *y*_*ij*_ is nonzero. *z*_*i*_ represents the probability of postsynaptic spiking after each presynaptic spike and we fit a smooth curve over the distribution of *z*_*i*_′s and their corresponding inter-spike intervals to compare with the empirical spike probability patterns.

**Fig. 2:**
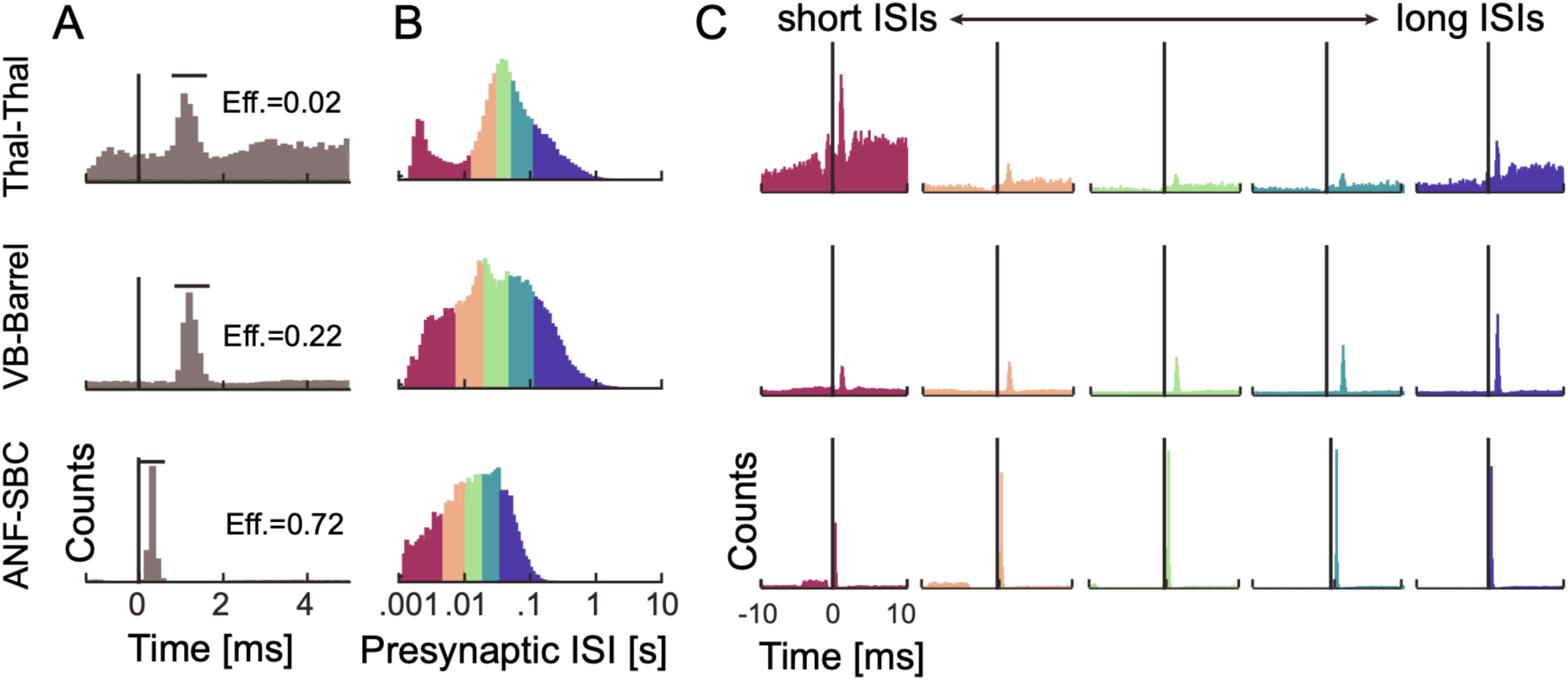
Spike transmission probability depends on the presynaptic ISI and differs between synapses. Cross-correlograms between pre- and postsynaptic spiking at three different synapses show an increase in the postsynaptic spike count (or probability) after a short latency, indicative of a monosynaptic connection. The efficacy (Eff.) for each synapse is calculated as the ratio between the number postsynaptic spikes that are above baseline in the transmission interval (denoted by the horizontal bar) and the number of presynaptic spikes. **B)** Inter-spike interval distributions (log-scale) for the presynaptic neurons. The distributions are color-coded into 5 quantiles with equal numbers of presynaptic spikes. **C)** We calculate a separate cross-correlogram using the subset of presynaptic spikes where the preceding spike fell within each ISI range. Colors correspond to (B) going from shorter presynaptic ISIs (left) to longer ISIs (right). Note that both the baseline firing rate and the synaptic peak for each connection change as a function of presynaptic ISI.

### Modeling the effect of local patterns of pre- and postsynaptic spiking

The observed and modeled spike transmission patterns, as calculated above, reflect the expected postsynaptic spike probability given a specific presynaptic ISI. However, since the presynaptic ISIs are not independent and there are serial correlations in ISIs, the detailed sequence of the pre- and postsynaptic spiking likely affects the shapes of these curves. To quantify the effects of serial ISI correlations on the model of spike transmission probability we demonstrate how local patterns of presynaptic spiking modifies spike transmission patterns in the data and the model. For each of the three strong identified synapses we measure postsynaptic spiking probability in response to presynaptic spike triplets. Due to the limited number of spikes in our data, we divide the presynaptic ISI distribution into few log-spaced intervals and measure the postsynaptic spiking probability for triplets with the two ISIs that fall in those intervals. Similarly, we measure the predicted postsynaptic probability in response to the presynaptic triplets. After measuring postsynaptic responses to presynaptic spike triplets in the data and the model, we simulate the contribution of STP in shaping the transmission pattern in response to these triplets. To factor out contributions of the postsynaptic history and slow changes in presynaptic firing rate, we fix the corresponding values in the model to their average values within the model. In these simulations, we also fix the initial values of the STP dynamics in the TM model for the first spike of the triplets to the average R and *u* within the model. This approach enables us to illustrate how short-term synaptic plasticity in triplets of presynaptic spikes changes spike transmission probability and how serial correlations in presynaptic spiking affect spike transmission probability.

The postsynaptic spike history and the serial correlations between the pre- and postsynaptic spiking also modify spike transmission probability patterns. To investigate history effects in the local pattern of pre- and postsynaptic spikes, we measured the postsynaptic spiking probability in response to two presynaptic spikes and a postsynaptic spike preceding the most recent presynaptic spike. Due to the limited number of spikes and sparseness of the split cross-correlograms, we again divided the presynaptic and postsynaptic ISI distributions into a few log-spaced intervals. We then measure the spike transmission probability for a group of presynaptic spikes that their preceding presynaptic ISIs and postsynaptic spike ISIs fall into different combinations of pre- and postsynaptic log-spaced intervals. After measuring postsynaptic responses to any possible combination of the two most recent presynaptic spikes and their postsynaptic spikes in the data and the model, we simulate the contribution of the history and STP together in shaping the transmission. In our simulation the excitability was set to the model estimates. To measure the effects of postsynaptic spiking history, for each postsynaptic ISI, we fix the history contribution to estimated post-spike history filter value at that postsynaptic ISI. We then use the predicted STP parameters from the data to simulate the STP contribution in response to paired pulses of presynaptic ISIs where we again fix the initial values of the TM model for the first presynaptic spike to the average R and *u* within the model. This approach enables us to illustrate how short-term synaptic plasticity in local patterns of two presynaptic spikes and a postsynaptic spike changes spike transmission probability and quantifies how serial correlations between pre- and postsynaptic spiking affect spike transmission probability.

### Evaluating prediction accuracy

In addition to evaluating the estimated parameters and comparing the model to empirical spike transmission probabilities, we also assess how accurately the model can predict postsynaptic spiking. Not only can we predict the probability of a spike given specific presynaptic ISIs, but we can also predict whether there will be a postsynaptic spike following each individual presynaptic spike. To quantify how well the predicted postsynaptic spike probability, *z*_*i*_, predicts the postsynaptic spiking activity, we use Receiver Operating Characteristic (ROC) curves. To compute the ROC curve, we first create a threshold version of *z*_*i*_ which operates as our prediction: 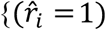 if (*z*_*i*_ > thr); 0 otherwise}. Changing the threshold from 0 to 1 traces out a relationship between the true positive rate (TPR) and false positive rate (FPR). The area under the ROC curve (AUC) reflects the performance of each model, where a perfect classifier has AUC=1 and a random classifier has AUC=0.5. Effectively, the AUC is the probability of a randomly chosen spike having a higher model probability than a randomly chosen non-spike (Hatsopoulos et al., 2007). Here we calculate the AUC for short intervals (∼5ms) after presynaptic spikes and check whether we detect a postsynaptic spike in the transmission interval where *α*(*t*) is above 10% of its maximum. Here we compare the AUC for the static model of connectivity without short-term synaptic plasticity with our dynamical model.

### A simplified rate model to simulate effects of synaptic summation and post-spike history

Our TM-GLM’s prediction of the spike transmission pattern is data-driven and depends on the full history of pre- and postsynaptic spiking. To better understand and illustrate how STP, synaptic summation, and post-spike history interact to create the observed patterns of spike transmission, we simulated postsynaptic responses in a simplified voltage model. Namely, we consider PSP summation in response to a pattern of two presynaptic spikes. We assume that the synapse is initially fully recovered, and the PSC amplitudes are determined by the 4-paramter TM model with *U* = 0.7, *τ*_*d*_ = 1.7, *τ*_*f*_ = 0.02, *f* =0.05 for the depressing synapse and *U* =0.1, *τ*_*d*_ = 0.02, *τ*_*f*_ = 1, *f* =0.11 for the facilitating synapse (Ghanbari et al., 2017). We then convolve the PSCs (delta function kernel) with a PSP kernel, exp(−t/*τ*_*ν*_) − exp (−t/*τ*_*r*_), with *τ*_*ν*_= .01 and *τ*_*r*_=.001 ms to describe synaptic summation. We assume that the instantaneous postsynaptic spike probability is simply a nonlinear function of the distance to a threshold voltage *σ*(5(*V*(*t*) – *V*_*th*_)) where *σ*(*x*) = 1/(1 + *e*−*x*) and *V*_*th*_ = .5, .75, and 1 correspond to strong, moderate, and weak inputs respectively. The spike transmission probability sums this instantaneous probability over a window of 20ms after each presynaptic spike. Finally, we adjust the spike transmission probability for the second PSP to account for potential post-spike history effects. Namely, we assume that the adjusted spike transmission probability for the second spike is 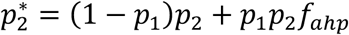 where *p*_1_ is the transmission probability for the first spike, *p*_2_ is the unadjusted probability for the second spike, and *f*_*ahp*_ is the effect of the after-hyperpolarization. Here we use *f*_*ahp*_(Δ*t*) = (*σ*(150(Δ*t* − 0.02)) − *c*)/*d* where Δ*t* is the presynaptic ISI, and *c* and *d* are constants ensuring that *f*_*ahp*_(0) = 0 and *f*_*ahp*_(∞) = 1. Although this simulation is highly simplified, it demonstrates how the observed spike transmission pattern depends, not just on the type and timescale of STP, but on the interaction between STP, synaptic summation, after-hyperpolarization effects, and the spike nonlinearity.

### Simulation of non-connections

The TM-GLM relies on correctly identifying monosynaptic connections. To investigate how our model performs when there is no actual synapse, we simulated a microcircuit with three neurons where a presynaptic neuron provides excitatory input to two postsynaptic neurons with different delays (1 and 3 ms). Here we test how different combinations of STP (depression and facilitation) in connections between pre- and postsynaptic neurons would impact the overall estimation of spike “transmission” probability in the spurious connection between the two postsynaptic neurons. Here the spikes of the presynaptic neuron were simulated from an inhomogeneous Poisson process with random, smooth rate fluctuations (5Hz average, 4.6Hz sd). The postsynaptic neurons were then simulated using a leaky integrate-and-fire neuron with spike frequency adaptation (parameters are from (Ghanbari et al., 2017)) that received a white noise current as input as well as a current-based synapse from the presynaptic neuron (double exponential with rise time 1ms, decay time 10ms). The PSCs of the input then vary according to the Tsodyks-Markram model (parameters for depression/facilitation are as in (Ghanbari et al., 2017)).

## Results

Short-term synaptic plasticity directly affects synaptic information processing by altering the amplitude of presynaptic currents (Abbott and Regehr, 2004). However, in most neural systems it remains unclear how these presynaptic effects translate to modified postsynaptic spike probability. Postsynaptic spiking is affected by many factors including short-term plasticity, postsynaptic spike history, summation of PSPs, and slow fluctuations in excitability. Here we develop a statistical model that includes each of these factors and allows their effects to be estimated solely using pre- and postsynaptic spiking activity. We examined the model’s ability to capture the observed patterns of spike transmission probability for three strong putative or identified synapses. We then use one of these systems (the endbulb of Held synapse in the auditory brainstem), to explore how the short-term dynamics of spike transmission depend on an external stimulus and compare the results with a model without short-term synaptic plasticity. Finally, we apply our model to spiking data from a large-scale, multi-electrode array recorded from multiple areas in an awake mouse. Here we investigate the STP dynamics in putative synapses from excitatory neurons onto two putative inhibitory neuron subtypes. We find that these two types of connections have distinct patterns of spike transmission, consistent with previous experimental observations.

### Spike transmission probability varies strongly as a function of presynaptic ISIs

Cross-correlograms of excitatory monosynaptic connections show a rapid, transient increase in the postsynaptic spiking probability shortly after the presynaptic spike, with a latency of ∼2-4ms (Perkel et al., 1967; Fetz and Gustafsson, 1983; Fetz et al., 1991; Poliakov et al., 1996). The timing and shape of the cross-correlogram depends on the presynaptic axonal conduction delay, the synaptic delay, and the strength of the connection. However, in the overall cross-correlogram the effects of all presynaptic spikes are averaged and any variations in spike transmission, such as dependence on the history of presynaptic spiking, are hidden (Fig. 2A). To quantify how the history of presynaptic spiking influences spike transmission probability, the probability of observing a postsynaptic spike shortly after a presynaptic spike, previous studies have compared the cross-correlograms for specific subsets of presynaptic spikes. For instance, comparing the cross-correlograms calculated for presynaptic spikes within defined inter-spike intervals (ISI) demonstrates how spike transmission probability varies depending on recent presynaptic spiking (Swadlow and Gusev, 2001; English et al., 2017). Here, to illustrate the diversity of short-term dynamics in spike transmission, we examine three strong synapses from three distinct neural systems: (i) a pair of neurons in thalamus in a male mouse, (ii) a projection from ventrobasal thalamus to somatosensory barrel cortex (VB-Barrel) in a female rabbit, and (iii) the auditory nerve fiber to spherical bushy cell projection in a female gerbil (ANF-SBC), the endbulb of Held. The short-term synaptic dynamics of thalamocortical projections, have been extensively characterized *in vivo* (Swadlow and Gusev, 2001; Stoelzel et al., 2008, 2009). Similarly, ANF-SBC synapses have been extensively studied in previous experiments and are well-characterized *in vitro* (Thomson et al., 2002; Yang and Xu-Friedman, 2008, 2009). The presynaptic neurons in each of these pairs have distinct ISI distributions (Fig. 2B), and, after splitting the spikes into ISI quantiles and calculating the correlogram for each quantile, we find that postsynaptic responses differ following short and long presynaptic ISIs (Fig. 2C). For the pair of thalamic neurons, spike transmission probability is increased at short and long intervals and reduced for mid-range ISIs (based on n=62661 presynaptic spikes). For the VB-Barrel connection, transmission probability is higher for longer ISIs (based on n=68345 presynaptic spikes), while for ANF-SBC the highest transmission probability occurs at intermediate intervals (based on n=20547 presynaptic spikes). These three cases illustrate that the short-term dynamics of spike transmission can be highly diverse between neurons and brain regions.

### The shape of spike transmission patterns depends on multiple factors

One potential explanation for the diverse dynamics of short-term spike transmission (Fig. 2) may be that some synapses are depressing while others are facilitating. Short-term synaptic plasticity directly alters postsynaptic currents such that the response after each presynaptic spike depends on the recent history of presynaptic spiking (Markram et al., 1998; Ghanbari et al., 2017). However, many factors can influence spike timing in addition to the dynamics of a single synapse. At short presynaptic ISIs, membrane potential summation can lead to larger PSPs and increased spike probability, even in absence of short-term synaptic plasticity (Carandini et al., 2007). Additionally, the spiking nonlinearity and the history of postsynaptic spiking can alter how a given pattern of presynaptic input is transformed into postsynaptic spiking (Pillow et al., 2008; Huang et al., 2016). To illustrate how STP, synaptic summation, and postsynaptic history interact to create a particular spike transmission pattern we performed simulations using a simplified spiking model with linear voltage summation, short-term plasticity, a soft spiking nonlinearity, and an after-hyperpolarization (Fig. 3).

**Fig. 3:**
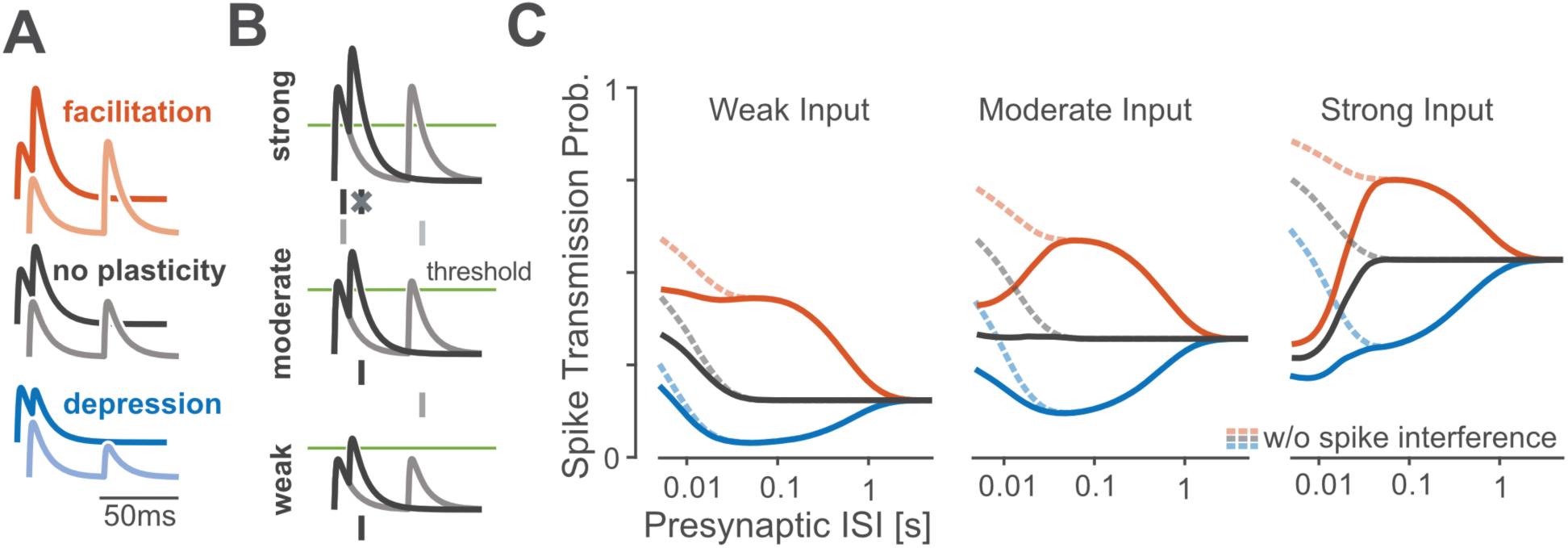
A simulation of a simplified spiking model shows how spike transmission probability depends on multiple factors. **A:** For different types of short-term synaptic plasticity, postsynaptic summation increases the amplitudes of the postsynaptic potentials (PSP) at shorter ISIs. Lines denote the membrane potential of a postsynaptic neuron in a simplified model as it responds to short (dark traces) and long (light) paired presynaptic pulses. Relative amplitudes of excitatory PSPs increase or decrease under the simplified model depending on the type of STP. **B:** Spike generation changes with synaptic strength. In this paired-pulse stimulation paradigm, stronger synapses are more likely to generate a spike following the first presynaptic impulse which can then decrease the spiking probability following the second impulse if there are post-spike history effects. As in (A) traces denote postsynaptic membrane potential responses to short (dark) and long (light) presynaptic ISIs. Dashes denote example postsynaptic spiking, with “spike interference” occurring for strong synapses and short ISIs. **C:** The pattern of spike transmission probability under the simplified model changes depending on the type of STP, the coupling strength, and presence of post-spike interference. Dashed lines show transmission probability without interference from previous postsynaptic spikes, while solid lines show how post-spike history effects can decrease the spike transmission probability.

Similar to experimental data (Markram et al., 1998; Ghanbari et al., 2017), the spike transmission probability in this simplified model depends on the presynaptic ISI as well as the type of STP. For depressing synapses, the spike transmission probability increases for longer presynaptic ISIs while for facilitating synapses it increases for mid-range ISIs. Independent of STP type, PSPs sum at short ISIs (Fig. 3A). However, in this model, the exact shape of transmission probabilities also depends on the strength of the synapse and the history of postsynaptic spiking. An after-hyperpolarization current following each postsynaptic spike, for instance, can briefly decrease the probability of subsequent spikes. In our simulation, we find that “spike interference” from previous postsynaptic activity can counteract membrane potential summation (Fig. 3B). This type of postsynaptic spike interference generally decreases the spike probability for shorter presynaptic ISIs, but the magnitude of this decrease depends on the synaptic strength and type of STP (Fig. 3C). Together, these simulations illustrate how patterns of spike transmission probability are the result of, not just STP, but of the complex interaction between the membrane potential, the spike nonlinearity, the post-spike history, and short-term synaptic plasticity.

### Spike transmission patterns are diverse across regions and species

The combination of synaptic and non-synaptic factors could be one explanation for the diversity of spike transmission patterns in experimental data. Here we aim to model these contributions and extend a previously developed generalized linear model (GLM) framework for static functional connections (Harris et al., 2003; Truccolo et al., 2005; Pillow et al., 2008). In the previous, static GLM the probability of postsynaptic spiking is modeled as a linear combination of a baseline firing rate parameter, a post-spike history filter to capture the postsynaptic spike dynamics, such as refractoriness and burstiness, and a coupling filter describing the fixed influence of presynaptic spikes. The sum of these effects is then passed through a spiking non-linearity. In our extended model we added a linear term that allows changes in the excitability of the postsynaptic neuron as a function of time (timescale >1 min) and allow the coupling term to change for each presynaptic spike according to the Tsodyks and Markram (TM) model of STP (Markram et al., 1998). We fit the parameters of this TM-GLM using only the pre- and postsynaptic spike observations and obtain parameters for each effect using approximate maximum likelihood estimation (see Methods). This provides estimates of the history and coupling filters, as in a static GLM, as well as additional parameters for the dynamical synapse (TM model), including facilitation, depression, membrane time-constants, and release probability. Given these parameters, this TM-GLM model provides estimates of the postsynaptic spiking probability following each observed presynaptic spike and can also predict spike transmission probabilities in response to arbitrary patterns of presynaptic inputs.

After fitting the model to pre- and postsynaptic spike-trains, we compared its behavior to experimentally observed patterns of spike transmission probability. In particular, we compare peaks in the split cross-correlograms to the average model prediction for the same sets of presynaptic spikes (see Methods). We find that our model is flexible enough to explain the changes in spiking transmission probability observed in spiking statistics for all three synapses above (Fig. 4A). Moreover, using the model-based approach, the contributions of the synaptic and non-synaptic component can be disentangled. Our results suggest that the pattern of spike transmission probability for the thalamus connection is dominated by a combination of membrane potential summation and short-term depression. Although depression decreases spike transmission probability at shorter ISIs, membrane summation acts to increase postsynaptic spiking. The ANF-SBC synapse, in contrast, shows an increase in spike transmission probability for a medium range of ISIs that is explained by a model dominated by short-term facilitation. Lastly, the VB-Barrel connection shows a higher postsynaptic response for spikes following longer ISIs (isolated) that is explained by the model as an effect of short-term synaptic depression.

**Fig. 4:**
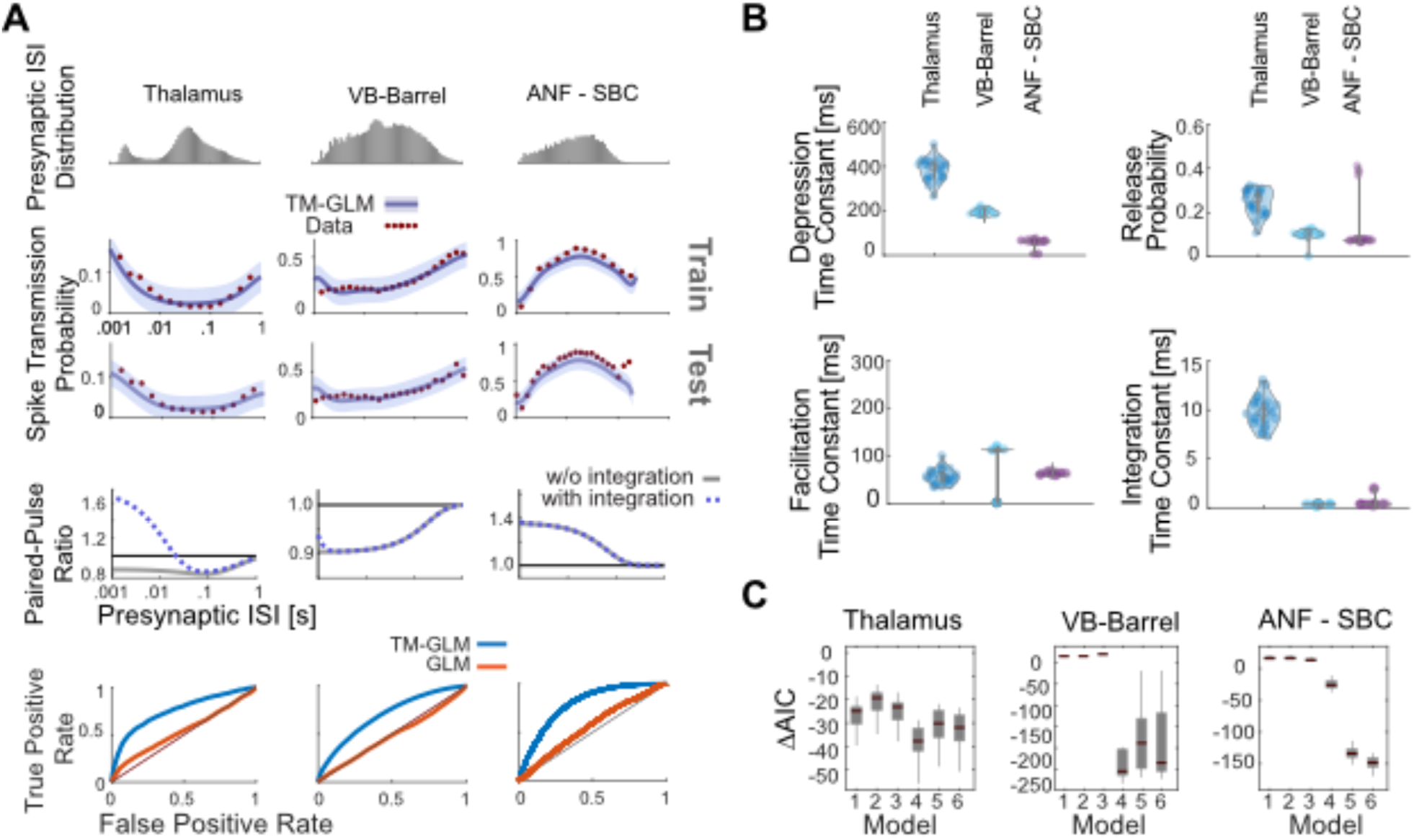
Including short-term dynamics substantially improves the model of spike transmission. **A:** Spike transmission patterns are diverse across different connections. For three different connections (between a pair of neurons in thalamus, a projection from ventrobasal thalamus to somatosensory cortex, and an auditory nerve fiber projection onto a spherical bushy cell) transmission patterns are modeled by a combination of different factors. For each synapse, top panels show the presynaptic ISI distributions (log-spaced). In the second/third row, the observed spike transmission probability (red data points) and model predictions (blue with 95% confidence bands) for training and test set (2-fold cross-validation). We then used the estimated TM parameters for each synapse and simulated responses to paired presynaptic pulses. Blue curves denote the PPRs of the full model, and gray lines denote PPRs by taking synaptic summation out. (bottom row) TM-GLM (blue) are superior in predicting individual postsynaptic transmission events compared to GLM (orange, without STP) for each synapse type. For each individual presynaptic spike, we compare the model transmission probability with the observed binary outcome. ROC curves show the prediction accuracy with positive deviations from the diagonal indicating better performance. **B:** Estimates for the four STP parameters of the model for each synapse. Dots represent estimates from bootstrap sampled data. **C:** Model comparison for 6 different models (Akaike information criteria, AIC, relative to a model without plasticity). Models: 1) Integration only, 2) Facilitation only, 3) Depression only, 4) 3-parameter TM, 5) 4-parameter TM without resetting integration, and 6) Full model. Boxplots denote the difference in AIC values for bootstrap samples in (B).

In addition to estimating the contributions of synaptic and non-synaptic factors that affect spike transmission, the model also improves the prediction of postsynaptic spiking. Although the cross-correlogram provides an average efficacy for spike transmission, our models provide detailed predictions of the postsynaptic spike probability following each presynaptic spike. Here we measure the Receiver Operating Characteristics (ROC curves) of our models during this short window of time following a presynaptic spike (see Methods). We compare the prediction of postsynaptic spiking activity in the full, dynamic synapse model and a static synapse model containing all components except STP. In all three datasets, a model with short-term synaptic plasticity provides substantially better predictions of the postsynaptic spiking activity. For the model with short-term synaptic plasticity accuracies were AUC=0.75±0.005, 0.69±0.002, and 0.79 ± 0.011 (mean ± SE) for the Thalamus pair, VB-Barrel, and ANF-SBC connections, respectively; compared to a model without STP where the model accuracies were AUC=0.54±0.003, 0.48±0.002, and 0.56±0.003 (mean±SE, bootstrapping over presynaptic spikes). Note that, although static synapse models do account for the average increased probability of spiking following a presynaptic spike, the fact that the AUC values are near chance (0.5) indicates that they do not accurately predict which presynaptic spikes will lead to a postsynaptic response and which will not.

In our model, the short-term dynamics of spike transmission are described by two coupled differential equations with five parameters: *θ*_*stp*_ = {*τ*_*d*_, *τ*_*f*_, *U, f, τ*_*s*_} (see Methods). Here we estimate values for depression, facilitation, and membrane time-constants along with release probability, *U*, and magnitude of facilitation, *f*, (Fig. 4B). Since these values are estimated from spikes and in observational settings rather than controlled experiments, the parameter estimates are likely to be biased by omitted variables (Stevenson, 2018). However, the parameter estimates do provide accurate predictions of postsynaptic spiking during natural, ongoing pre- and post-synaptic spiking, and may provide an initial, approximate description of synaptic dynamics. Comparing the estimates for the three model synapses – the thalamus pair has the highest release probability (0.29±0.04 SE) and the largest membrane (14±2 ms) and depression time-constants (410±107 ms). The VB-Barrel connection has a small membrane time-constant (0.3±0.003 ms) and a larger depression (182±8 ms) time-constant than facilitation time-constant (105±9 ms). The ANF-SBC synapse has the lowest release probability of the three connections (0.068±0.006) and small depression (67±6 ms) and membrane time-constant (0.25±0.02 ms). Due to the potential for omitted variable bias and differences in experimental preparations comparing these values directly to measurements from intracellular recordings is difficult. However, the values estimated from ongoing spiking and the results from intracellular recordings are generally in agreement. For instance, previous *in vitro* studies of thalamocortical projections found that paired-pulse ratios ranged from 0.3-0.9 consistent with depressing VB-Barrel synapses (Gil et al., 1997). Additionally, *in vitro* observations of ANF-SBC connections report depression time-constants on the order of 2-25 ms in response to a 100 Hz stimulus train (Wang and Manis, 2005, 2008). These previous estimates are substantially faster than the time-constants estimated by the TM-GLM for the ANF-SBC connection here. However, different patterns of presynaptic input (e.g. regular, Poisson, natural) or differences in calcium concentration and temperature may make it difficult to compare *in vitro* and *in vivo* STP parameters directly. One parameter that may be more readily comparable across preparations is the membrane time-constant. We find that the estimated membrane time-constant from the TM-GLM for the thalamus pair is consistent with thalamus relay cells observed intracellularly (12.2 ± 1.1 ms, n=8) (Paz et al., 2007), and the estimated membrane time-constant for ANF-SBC is close to *in vitro* measurements (1.05 ± 0.09 ms) as well (Wang and Manis, 2005).

The TM-model used here is one of many possible parametric descriptions of short-term plasticity (Hennig, 2013). Previous work modeling intracellular recordings suggests that the full TM model may not be necessary to explain STP at some, purely depressing synapses (Costa et al., 2013). Therefore, we explored how simplified TM models of STP, with fewer parameters, compare with the full model using the Akaike information criterion (AIC; see Methods and Fig 4C). AIC evaluates model accuracy (log-likelihood) penalized by the number of parameters, and lower AIC may indicate that a simplified model with fewer parameters is preferred over a more complex model. Generally, the synaptic dynamics in this class of models can be described by four parameters: a time-constant for depression *τ*_*d*_, a time-constant for facilitation *τ*_*f*_, a baseline release probability *U*, and facilitation parameter *f*. When modeling spike transmission we additionally include a parameter for the membrane time-constant *τ*_*s*_ and consider the possibility that the membrane potential “resets” following a post-synaptic spike (see Methods). For each of these models, it is important to note there may be many possible parameter settings that are consistent with the data, particularly when the recording time is limited (Costa et al., 2013). These redundancies are present even in simple quantal analysis methods (Bykowska et al., 2019). Here, altogether, we compare our full model to five reduced models: 1) a model with only membrane integration, without dynamic release probability and resources, 2) a facilitation only model, 3) a depression only model, 4) a 3-parameter TM model where the magnitude of facilitation is fixed, and 5) the full TM model, but without post-spike reset of integration (Table 1). The full TM model performs competitively in all cases, but, for some synapses, just as with previous results modeling PSPs (Costa et al., 2013), the full model may be overly flexible and simpler models, with fewer parameters, may be preferred. For the thalamus pair and VB-Barrel projection, the 3-parameter TM-model with fixed magnitude of facilitation has the lowest AIC (p<10^−9^ and p=0.07 compared to model 6 with a paired t-test). For the ANF-SBC connection the full model gives the lowest AIC (p<10^−6^ compared to model 4). For all three connections, models 4-6 perform statistically significantly better than both the model without STP (e.g. Δ*AIC*<0, Bonferroni-corrected paired t-test p<0.001) and model 1 (Bonferroni-corrected paired t-test, p<0.001). These results provide further evidence for STP-like changes in spike transmission at these connections.

**Table 1:** Parameters included each model. Note that *τ*_*s*_ is not constrained in any of the 6 models.

| Model | Description | $\tau_d$ | $\tau_f$ | $f$ | $U$ | Reset |
| --- | --- | --- | --- | --- | --- | --- |
| 1 | Integration only | 0 | 0 | 1 | 1 | Yes |
| 2 | Facilitation only | 0 | No constraints |  |  | Yes |
| 3 | Depression only | No constraints | 0 | No constraints |  | Yes |
| 4 | 3-parameter TM | No constraints | | $f = U$ | | Yes |
| 5 | TM without reset | No constraints |  |  |  | No |
| 6 | Full model | No constraints |  |  |  | Yes |

**Table 2:** Summary of parameter estimates from the full TM-GLM. Sample size (*n*), membrane time-constant (*τ*_*s*_), depression time-constant (*τ*_*d*_), and facilitation time-constant (*τ*_*f*_), and release probabilities (*U*) for the identified and putative synapses from our three case studies and multi-electrode recordings. For the cases studies, the mean±standard deviation is shown for the bootstrap samples. For the MEA data, the mean±sd is shown across putative connections. In all cases, the parameters are estimated from ongoing, in vivo spiking activity.

| Synapse | $n$ | $\tau_s$ (ms) | $\tau_d$ (ms) | $\tau_f$ (ms) | $U$ |
| --- | --- | --- | --- | --- | --- |
| Thalamus | 1 | $14 \pm 2$ | $410 \pm 107$ | $37 \pm 12$ | $0.29 \pm 0.04$ |
| VB-Barrel | 1 | $0.3 \pm 0.003$ | $182 \pm 8$ | $105 \pm 9$ | $0.10 \pm 0.05$ |
| ANF-SBC | 1 | $0.25 \pm 0.02$ | $67 \pm 6$ | $71 \pm 3$ | $0.068 \pm 0.006$ |
| excitatory-RS | 19 | $84 \pm 116$ | $215 \pm 219$ | $820 \pm 745$ | $0.46 \pm 0.17$ |
| excitatory-FS | 22 | $72 \pm 196$ | $411 \pm 459$ | $406 \pm 552$ | $0.34 \pm 0.19$ |

### Recent patterns of pre- and postsynaptic spiking shape the synaptic transmission probability

Although previous studies have focused largely on how spike transmission probability varies as a function of the single ISI preceding the most recent presynaptic, synaptic dynamics depend on the full sequence of presynaptic spiking. Unlike *in vitro* experiments where the state of the synapse can, to some extent, be controlled before studying responses to a specific presynaptic pattern, *in vivo* measurements of spike transmission can be heavily influenced by higher-order correlations between successive ISIs (Stoelzel et al., 2008). Additionally, it is difficult to assess the effects of multi-spike patterns empirically by splitting the correlograms, since the number of observations for any given presynaptic spike pattern rapidly decreases with the number of spikes in the pattern. Here we examine how spike transmission depends, not just on the preceding presynaptic ISI, but on triplets of spikes. We compare the empirically observed spike transmission probability following triplets to the estimated spike transmission probability from the TM-GLM. Using the model fits for TM-GLM, we then simulate postsynaptic responses to isolated patterns of spikes and determine to what extent the observed spike transmission patterns are influenced by higher-order correlations between successive ISIs.

First, in addition to the timing of the two preceding presynaptic spikes (separated by the interval ISI_1_), we split correlograms based on the timing of the three preceding presynaptic spikes (Fig. 5A), separated by the most recent interval and the one before (ISI_2_). Since the TM-GLM provides estimates of the post-synaptic spike probability following every presynaptic spike, we can split both the data and model fits the same way (Fig. 5C). We find that the spike transmission patterns clearly depend on the triplet patterns of presynaptic spikes in ongoing spiking activity. That is, the spike transmission probability is influenced by both ISI_1_ and ISI_2_, and the interaction between the two ISIs differs between synapses. However, as with spike transmission as a function of ISI_1_ alone, the TM-GLM accurately captures the patterns of spike transmission for triplets of presynaptic spikes for the three synapses. In the thalamus pair, spike transmission probability is most influenced by ISI_1_, and the effect of ISI_2_ appears to be weak or, at least, does not appear to be monotonic. Spike transmission probability at the VB-Barrel connection depends on both ISI_1_ and ISI_2_, with higher spike transmission probability for longer ISI_2_, consistent with recovery from depression. Lastly, for the ANF-SBC connection, transmission probabilities decrease for shorter ISI_2_, but there also appears to be a strong interaction between ISI_1_ and ISI_2_, where transmission probability is high for multiple combinations of these two intervals (e.g. intervals of 10 ms then 100 ms and intervals of 100 ms then 10 ms both result in high probability transmission).

**Fig. 5:**
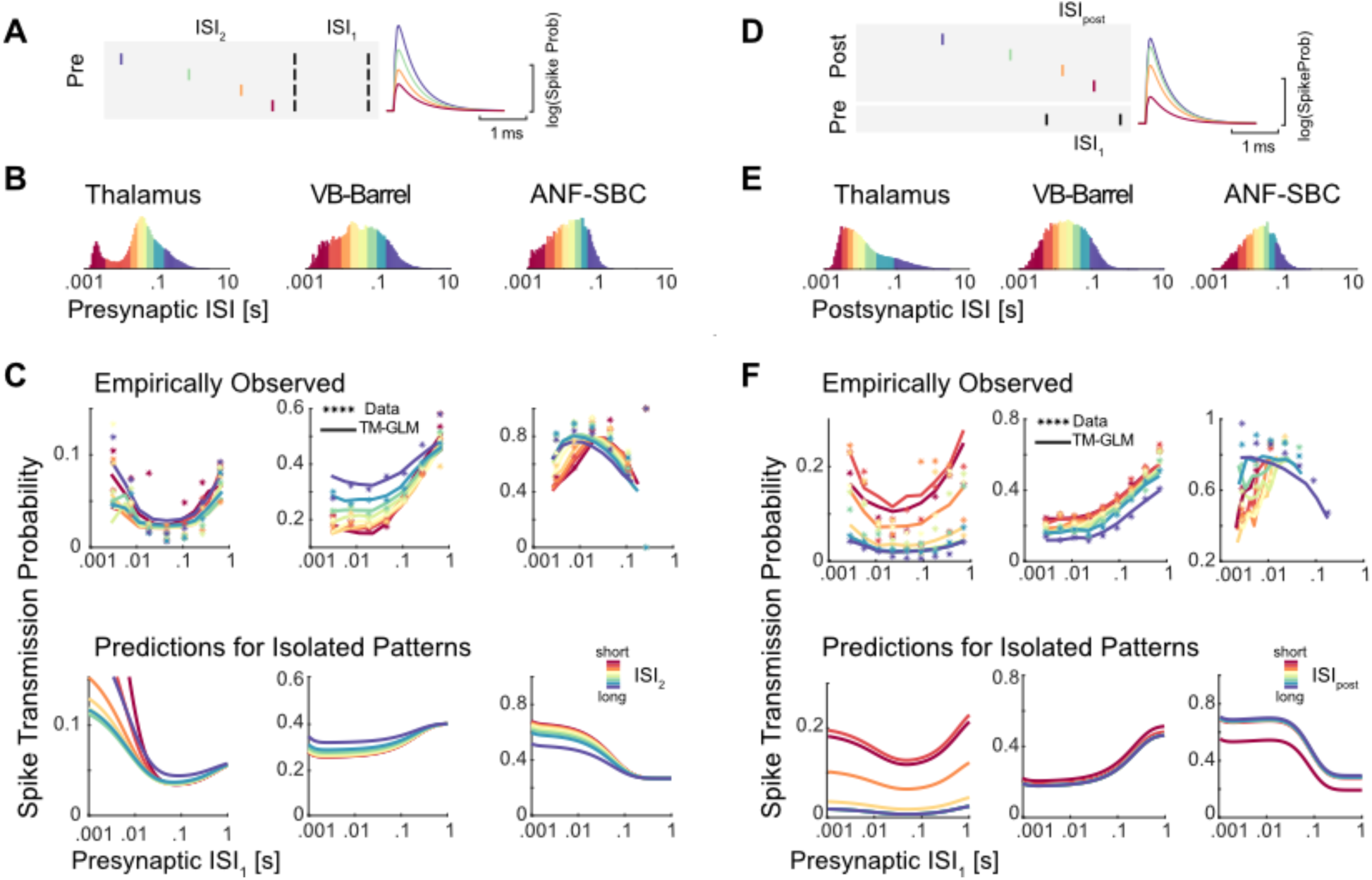
Pre- and postsynaptic spiking history determine transmission probability. A) Schematic of 4 different patterns of presynaptic spike triplets with a fixed interval between the two most recent presynaptic spikes (spikes denoted by black lines separated by ISI_1_). B) We then split the presynaptic ISI distribution into 8 quantiles, denoted by the different colors. C) We then assess how ISI_2_ influences the spike transmission previously described for ISI_1_. Using the natural occurrence of different ISI_1_ and ISI_2_ in the data, each data point shows the observed spike transmission probability for each pattern (colors correspond to ISI_2_ quantiles). Lines denote the average estimated transmission probability for each pattern under the model (based on the natural sequence of observed spikes). To examine the influence of serial correlations, we then simulate model responses to the isolated triplet pattern, assuming the synapse is initially in an average state (bottom panels). D) Synaptic transmission patterns change depending on the history of postsynaptic spiking, as well. E) Note that the postsynaptic ISI distributions need not match the presynaptic distributions. F) Here each data point in the scatter plots shows the spike transmission probability following different combinations of ISI_1_ and ISI_post_. Here, colors denote quantiles of the postsynaptic ISI distribution. Solid lines show the estimated transmission probability for each pattern under the model (based on the natural sequence of observed spikes). The bottom panels show model responses to isolated patterns using the estimated STP parameters and fixing the excitability from the model fits to their average values.

Although these empirical results suggest that spike transmission probability is influenced by triplet patterns of presynaptic spikes, these triplets are not isolated events but are embedded in longer sequences of spikes with higher-order correlations between successive ISIs. To examine to what extent the model predictions are affected by higher-order correlations between successive ISIs, we again use the estimated parameters in the TM-GLM to simulate postsynaptic responses to hypothetical, isolated triplets of presynaptic spikes (Fig. 5C, bottom). In these simulations we fix the post-spike history effect and the excitability in the model to their average values from model fits, and we fix the initial STP state (initial values of *R* and *u* in TM model) for the first spike in triplets to the average *R* and *u* values from the model fits. Although the initial states of the pre- and postsynaptic neurons in the experimental data are not matched for different values of ISI_1_ and ISI_2_, by simulating, we can assess the isolated influence of different triplets (ISI_1_ and ISI_2_) on the model. Here we find that for the thalamus pair, although the empirical data showed no clear effect for ISI_2_, the simulated spike transmission probability increases with short ISI_2_, consistent with strong synaptic summation. One reason that this effect may be masked in the empirical transmission probabilities is that post-spike history effects could act to decrease the probability of future postsynaptic spikes. For the VB-Barrel simulations, we find that short ISI_2_ decreases transmission probability, consistent with the empirical transmission patterns, although less pronounced. Serial correlations in the sequence of presynaptic spikes (such as long bursts) could act to accentuate the depression in the empirical observations beyond what we see with the simulated responses to isolated triplets. Finally, for the ANF-SBC, although the empirical transmission probability showed decreased transmission for short ISI_2_, the simulated responses to isolated patterns have increasing transmission at short ISI_2_ (due to synaptic summation). This difference is likely due to the post-spike history filter, which has been fixed for the simulations, but can have a large effect in the experimental data. Since the overall efficacy of this synapse is quite high (>0.7), is likely that a postsynaptic spike follows the first or second presynaptic spike which then influences the response to the third spike.

To better understand the effects of post-spike history, we examined how the postsynaptic spiking history changes the spike transmission patterns with a similar approach. In addition to splitting the correlograms based on ISI_1_, we also split based on the previous postsynaptic ISI, ISI_post_ (Fig. 5D). Here, as with the triplets of presynaptic spikes, we find that the spike transmission patterns depend on the triplet patterns of 2 pre- and 1 postsynaptic spike in data and that the TM-GLM accurately captures the patterns of spike transmission at our three synapses (Fig. 5F). Here, for both thalamus and VB-Barrel pairs, synaptic transmission probability decreases after a long postsynaptic ISI for all values of ISI_1_. In contrast, the ANF-SBC connection shows decreased transmission probability at short postsynaptic ISIs.

As with the triplets of presynaptic spikes, we then simulate (Fig. 5F, bottom) how patterns of 2 pre- and 1 postsynaptic spike change spike transmission probability when the neurons start from the same initial conditions (average values of excitability, post-spike history, *R* and *u*). For the thalamus and VB-Barrel pairs, the simulations of isolated patterns match the general trends of empirical spike transmission. However, for the VB-Barrel synapse, the effect of ISI_post_ in the empirical transmission patterns is stronger than in the simulations, suggesting that serial correlations in ISIs could again play a role and amplify the effects of isolated patterns.

### Spike transmission patterns change depending on stimulus type

The results above suggest that the presynaptic spike pattern has a complex effect on spike transmission probability. In sensory systems, one factor that affects the presynaptic spike pattern is the external stimulus. To examine how differences in stimulus statistics might alter spike transmission, we fitted our model to a dataset recorded juxtacellularly from an ANF-SBC synapse, presented with *natural sounds*, a range of randomized frequency-level pure-tones (*tuning stimuli*), and s*pontaneous activity* in the absence of acoustic stimulation. Note that this dataset was partially (tuning stimuli) used in the first section of the results. We merged these three datasets and fitted the model to the merged dataset. As with the previous fits of the ANF-SBC connection (based on a different set of *tuning stimuli*), the transmission probability under all three conditions exhibits a bandpass-like pattern in mid-range ISIs suggesting facilitation and little to no synaptic summation. However, spike transmission during natural stimuli was markedly different from that during pure tone stimulation. During natural sounds, transmission probability is maximized at 100 ms rather than 10 ms found in the *tuning stimuli* and during *spontaneous activity*. Further, *natural stimuli* have much lower transmission probability at short ISIs. Interestingly, the TM-GLM captures the overall facilitation, but also captures differences due to the different stimuli (Fig 6A). In contrast, a static GLM captures almost none of the variations in spike transmission probability. Together, these results suggest that the combination of STP, synaptic summation, history, and excitability is sufficient to explain the observed differences spike transmission between stimuli, without requiring any additional adaptation or plasticity.

**Fig. 6:**
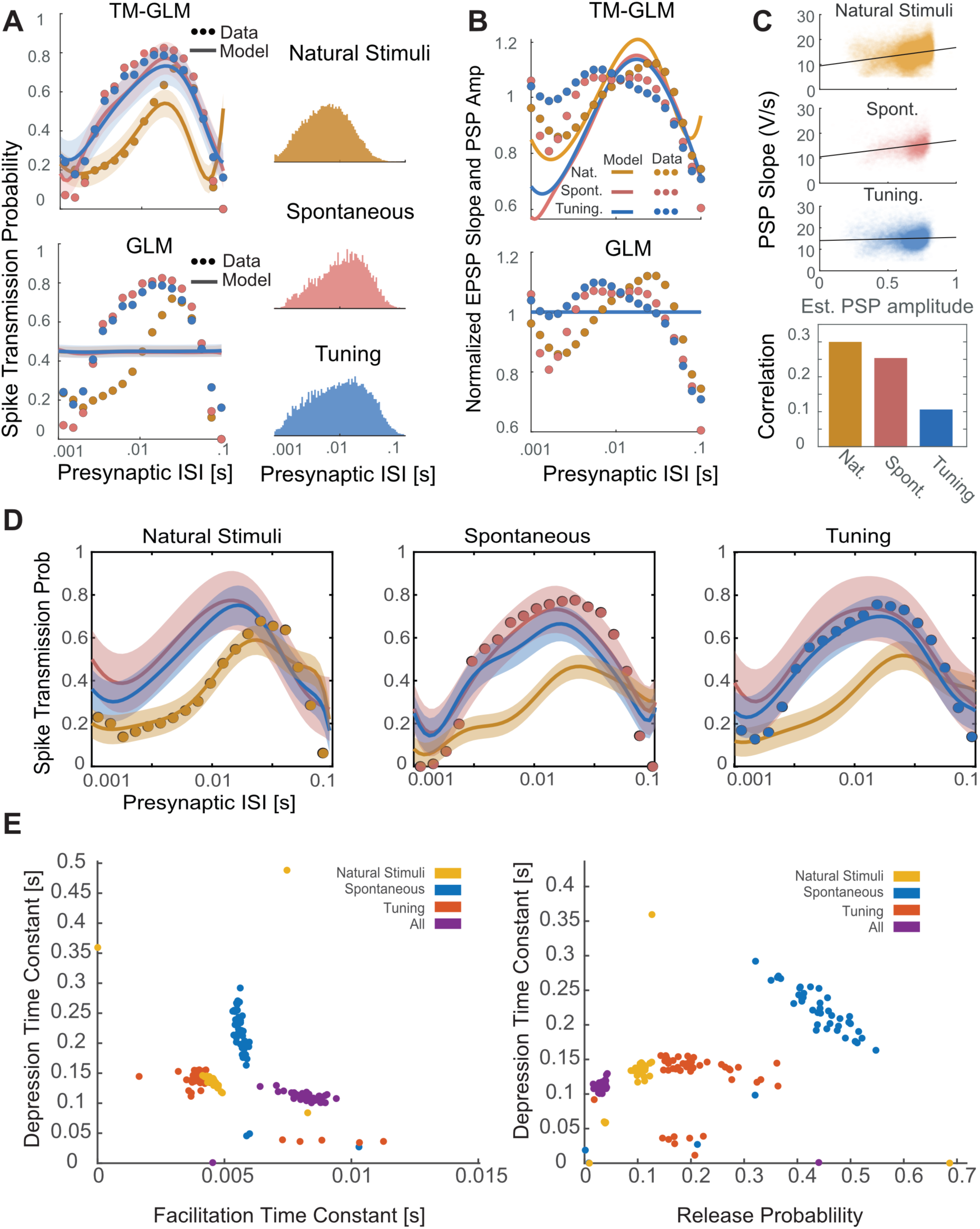
The TM-GLM captures stimulus-dependent changes in spike transmission probability at the ANF-SBC synapse. **A)** The TM-GLM captures stimulus-dependent spike transmission probability patterns better than a static model without short-term synaptic plasticity. Dots show spike transmission probability for (log-spaced) presynaptic ISIs during two types of auditory stimuli and during spontaneous activity: Natural Sounds (yellow), Spontaneous Activity (red), and Tuning Stimuli (blue). Solid lines and 95% confidence bands show model predictions for each stimulus type. Corresponding inter-spike interval distributions are shown on the right. **B)** The TM-GLM captures changes in extracellularly recorded PSPs. Here the observed PSP slope (dots) approximately matches the coupling term in the TM-GLM (solid lines) for each three stimuli. Although the spike transmission probability of the static GLM can vary as a function of presynaptic ISI due to non-synaptic factors, the coupling term is fixed. **C)** Estimates of individual PSP amplitudes predicted by the model and their PSP slopes in the juxtacellular recording. Black lines denote linear fits and the bar plot shows the corresponding Spearman correlations. **D)** After fitting each stimuli condition separately, in each column we plotted the estimated spike transmission probability of each type using the estimated STP parameters of others. **E)** Distribution of parameters from bootstrap samples with the TM-GLM fit for individual stimuli and all stimuli combined.

Since these recordings were performed juxtacellularly, we also have access to the slope of individual (extracellularly observed) PSPs, which are correlated with the intracellular PSP amplitudes. We compared patterns of individual PSP slopes for each stimulus type and examine how these slopes correlate with the estimated coupling amplitude following individual presynaptic spikes in our model (Fig. 6B, 6C). Note that patterns of PSP slopes do not have the same pattern as spike transmission probability, since there are other factors (e.g. postsynaptic spiking history) contributing to postsynaptic spiking. However, as with spike transmission, we find that the PSP amplitudes are stimulus-dependent and that a static GLM without STP cannot account for these variations. Additionally, although the correlation is not perfect, the individual coupling effects in the model do correlate with the measured PSP slope, even though the model is only fit to spikes. By modeling dynamic functional connectivity, we can approximately reconstruct the amplitude of individual synaptic events.

We then analyze how much the TM-GLM can generalize to other stimulus types when fit to one stimulus type. We find that, although the model can describe the spike transmission patterns for all three stimuli when fit to all stimuli, the model does not generalize to natural stimuli when fit exclusively to one of the other stimulus types (and vice versa, Fig. 6D). The parameters from each of these models are distinct – occupying different regions of the parameter space. Notably, the model fit to all stimuli has a lower release probability and a higher facilitation time-constant compared to the models fit to individual stimuli (Fig. 6E).

### Postsynaptic cell-type specific changes in spike transmission patterns

We also applied our model to spiking data from a large-scale multi-electrode array recording to investigate the spike transmission dynamics in synapses from putative excitatory neurons to two different putative inhibitory subtypes. We detected putative synapses using the log-likelihood ratio (LLR < −6, ∼200 synapses) between a full model of the correlogram that includes the synaptic effect and smooth model of the correlogram that only captures the slow structure (see Methods). We then found excitatory-inhibitory microcircuits where putative excitatory neurons (based on the cross-correlogram and spike waveform) give inputs to putative inhibitory neurons (41 excitatory synapses onto 9 inhibitory neurons in total). To identify inhibitory neurons as inhibitory, we required the neuron to have an outgoing connection to a third neuron with a fast, transient decrease in the cross-correlogram. Each of the 9 putative inhibitory neurons here had at least one outgoing connection where the spiking probability of a downstream neuron decreases >18% relative to baseline following its spiking (Fig. 7A). We then categorized each neuron as a putative fast-spiking (FS, n=5) or regular-spiking (RS, n=4) unit based on the spike waveform and firing rate (Fig. 7B). Putative FS units had narrow-width spike waveforms (half-width of the trough = 0.08±0.02 ms) and higher firing rates (26.07±9.6 Hz) compared to putative RS neurons (n=4) with broader waveforms (half-width = 0.14±0.02 ms) and lower firing rate (10.18±10.01 Hz).

**Fig. 7:**
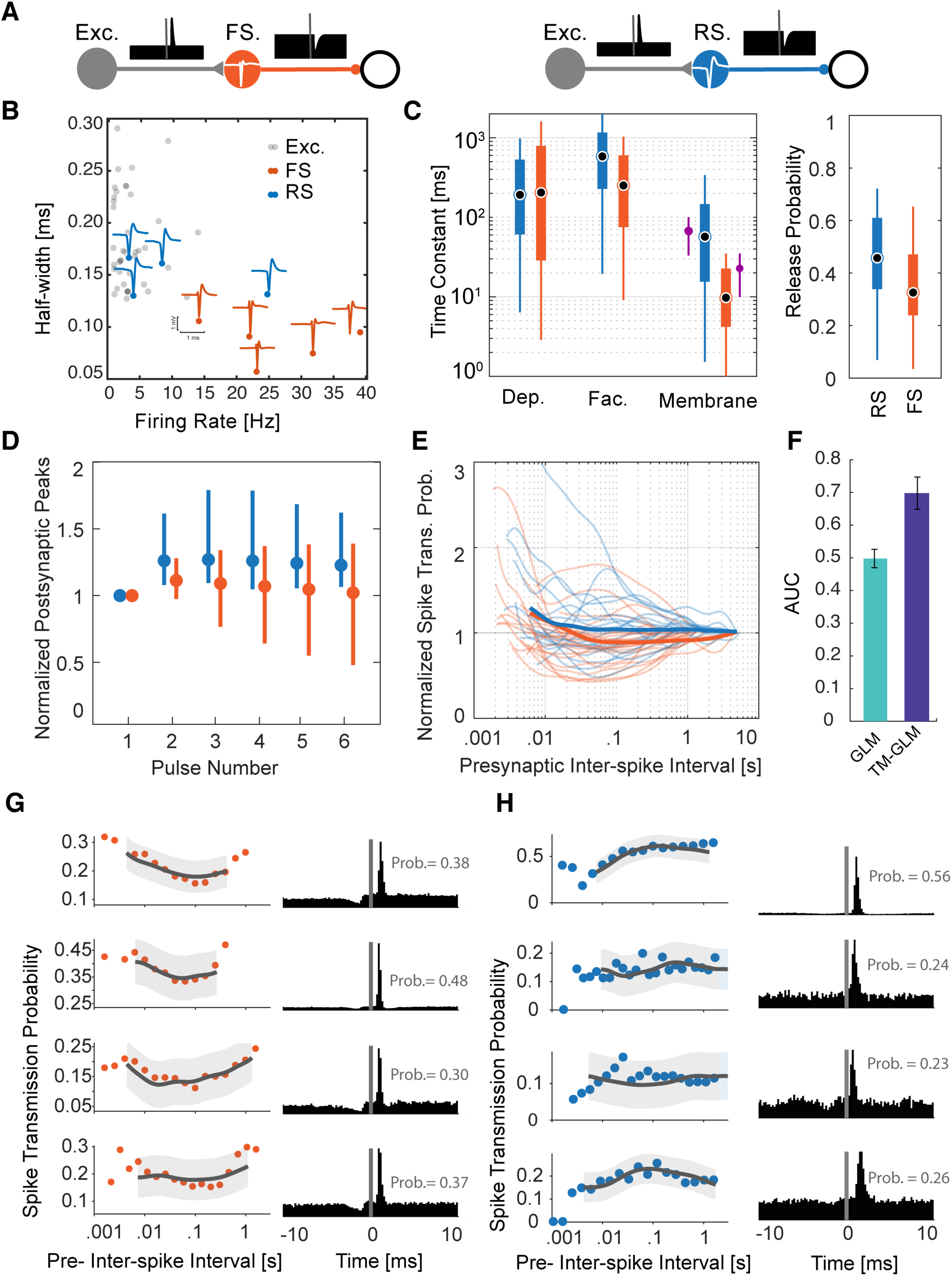
Distinctive short-term dynamics for spike transmission in connections between excitatory neurons to putative Regular-Spiking (RS) and Fast-Spiking (FS) inhibitory neurons. A) Here we examine putative synapses between excitatory neurons and inhibitory neurons (identified by their cross-correlations) and separate the putative inhibitory neurons into two classes: fast-spiking, which have narrow spike waveforms and high rates (left), and regular-spiking (right), which have wide waveforms and lower rates. Identifying these synapses requires both finding both a putative excitatory input and a putative inhibitory output for the same neuron. B) Half-widths (of the trough) of the spike waveforms and firing rates for the FS (orange) and RS (blue) inhibitory neurons, as well as, their excitatory inputs (grey). Individual blue and orange waveforms (maximum amplitude across the MEA) are shown for all 9 putative inhibitory neurons. C) Estimated depression, facilitation, and membrane time-constants for excitatory-RS and excitatory-FS connections, along with the release probability (right). The purple error-bar next to the membrane time-constant estimations show the median and standard deviations from *in vitro* experiments (Perrenoud et al., 2013). D) Simulated postsynaptic potential amplitudes estimated from Tsodyks-Markram model of short-term synaptic plasticity using estimated parameters. For each synapse, PSPs are estimated in response to a pulse train with inter-pulse intervals set to their corresponding average presynaptic inter-spike intervals. Dots and error bars denote the median and inter-quartile range for excitatory-RS (blue) and excitatory-FS (red) connections. These responses include the effect of membrane potential integration. E) Spike transmission probability patterns for individual synapses of excitatory-RS (blue) and excitatory-FS (red) connections normalized by long interval probabilities as a function of the presynaptic ISI. F) Area Under the Curve (AUC) of postsynaptic spiking prediction using the static GLM without short-term synaptic plasticity (green) and the TM-GLM with short-term synaptic plasticity (blue). G-H) Spike-transmission probabilities (left) and corresponding cross-correlograms (right) of 4 putative excitatory inputs to putative FS (G) and RS (H) inhibitory neurons show cell-type specific similarities.

We identified these microcircuits in different regions with 4 putative excitatory-inhibitory microcircuits recorded in hippocampus (depth differences: 77.2 ± 49.4 *μ* m), 3 in thalamus (49.4±26.2 *μ*m), and 2 in motor cortex (36.4±23.5 *μ*m). Putative excitatory neurons showed a wide spike waveform (half-width = 0.18 ± 0.04 ms) similar to the putative regular-spiking inhibitory neurons, but these two classes can be distinguished by their outgoing connection types (e.g. inhibitory/excitatory) (Moore and Wehr, 2013) (Fig. 7B). Average efficacies from putative excitatory-FS connections (0.22 ± 0.12, n=22) were larger, on average, compared to putative excitatory-RS efficacies (0.13±0.13, n=19). We then fit the TM-GLM to data from these 41 putative synapses, similar to the three identified synapses analyzed above. Again, due to omitted variable bias, the interpretation of the parameter values for the model fits is not necessarily straight-forward. However, we find that there is substantial overlap between the estimated STP parameters for excitatory connections onto these two inhibitory subtypes (Fig. 7C). The depression time-constant for excitatory-RS connections is 215± 219 ms (mean± SD, median 96 ms) and for excitatory-FS is 411±459 ms (median 191 ms). The facilitation time-constant for excitatory-RS connections is 820±745 ms (median 588 ms) and 406±552 ms (median 236 ms) for excitatory-FS connections. And the membrane time-constant for excitatory-RS connection is 84±116 ms compared to 72 ± 196 ms for excitatory-FS. Interestingly, the estimates for membrane time-constant (median 10 ms for FS, 45 ms for RS) are similar to the parameters measured using intracellular recordings *in vitro* (Perrenoud et al., 2013).

Previous *in vitro* studies of postsynaptic cell-type specific STP concluded that putative excitatory-RS connections show facilitation and putative excitatory-FS connections show depression (Thomson and Lamy, 2007). Moreover, few *in vivo* studies characterized stimulated activities in these connections (Pala and Petersen, 2015, 2018; Sedigh-Sarvestani and Vigeland, 2017). A cell-type-specific study of somatosensory connections *in vivo* using 50Hz optogenetic stimulation found little short-term plasticity in connections to Parvalbumin-expressing neurons (putative excitatory-FS here), while excitatory to Somatostatin-expressing neurons (putative excitatory-RS here) showed facilitation (Pala and Petersen, 2015). However, we are not aware of any *in vivo* experiments that measured depression or facilitation time-constants for these systems during ongoing spiking activity. Here we find that both connection types are somewhat facilitating but excitatory-FS connections having a slightly shorter facilitation time-constant. However, unlike what would be expected if excitatory-FS connections were depressing, the release probability of excitatory-FS connections is lower than excitatory-RS connections (Fig. 7C, 0.34±0.19 for FS, 0.46±0.17 for RS). To better understand synaptic transmission *in vivo* it is important to consider not just the parameters of the synapse but the full history of presynaptic spiking in the individual presynaptic neurons. We use the estimated model parameters to simulate responses to a train of regular presynaptic spikes with the frequency matched to the average firing rate of the corresponding excitatory input. In simulating postsynaptic responses to the spike train, we fix the excitability and postsynaptic history to their average values from model fits and set the initial STP state of the first spike in the train to the average *R* and *u* values from model fits. With these input-matched simulations, excitatory-RS connections show higher amplitude postsynaptic potentials compared to excitatory-FS connections (Fig. 7D, the effect of membrane potential integration is included). This is in accordance with the previously observed small degree of facilitation in connections to Somatostatin-expressing neurons and small degree of short-term plasticity in connections to Parvalbumin cells in (Pala and Petersen, 2015).

We also calculated spike transmission probabilities for all connections. On average, connections to regular-spiking inhibitory neurons show a higher spike transmission probability across interspike intervals (Fig. 7E). For all connections, we then evaluated the spike prediction accuracy of a model without STP (e.g. static GLM) with our TM-GLM using the Area Under the ROC Curve (Fig. 7F). The model with STP (TM-GLM) gives more accurate predictions for which presynaptic spikes will lead to postsynaptic spiking for our population of 41 putative excitatory-inhibitory connections (AUC=.69±.05) in comparison with the static GLM (AUC=.50±.03). Altogether, these results illustrate how a dynamic model of functional connectivity, such as the TM-GLM, can provide a detailed functional description of the short-term dynamics of spike transmission in awake, behaving animals.

### Spike “transmission” patterns between unconnected pairs of neurons

It is important to note that the dynamic functional connectivity model presented here assumes that, before fitting the model, we have accurately identified a monosynaptic connection. In some settings, it is possible to identify connections using optogenetic stimulation (English et al., 2017) or juxtacellular recording, however, in cases where we can only identify putative connections, it is important to consider the possibility that we are modeling a spurious correlation between neurons that are not actually monosynaptically connected. In general, the detection of monosynaptic connections from multielectrode spiking activity is far from perfect (Kobayashi et al., 2019).

To examine how the TM-GLM might be influenced by spurious correlations, we first simulated a small circuit with common drive that would likely lead to a falsely detected monosynaptic connection (Fig 8A). Here an unobserved presynaptic (inhomogeneous Poisson process) neuron provides strong excitatory input to two leaky integrate-and-fire postsynaptic neurons. Due to a difference in the latencies of these connections, there is a spurious peak in the correlogram between the two postsynaptic neurons where one postsynaptic neuron appears to excite the other. We find that when we measure the amplitude of this spurious peak, there are some variations as a function of the presumed presynaptic neuron’s ISI, and the spike “transmission” pattern varies depending on whether the projections from the true presynaptic are both depressing, both facilitating, or a mixture of depressing and facilitating (Fig 8B). However, the TM-GLM is nearly constant (∼0.1% variation) and does not accurately fit the observed variation. Despite a spurious correlation, the detailed pattern of spikes between the two postsynaptic neurons is unstructured and not well described by the TM model.

**Fig. 8:**
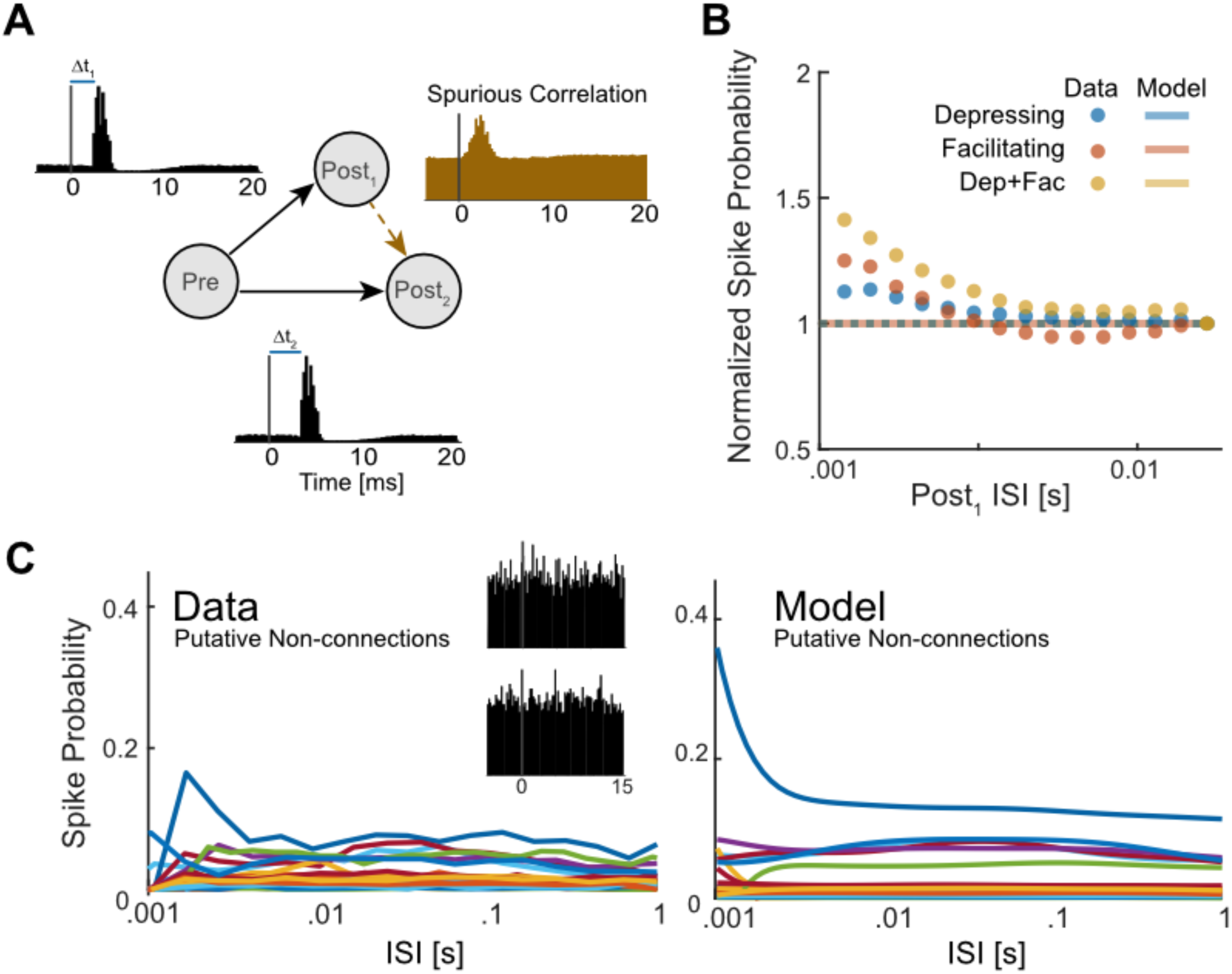
Short-term changes in spike probability for neurons that are not monosynaptically connected. A) In a simulated circuit, we generated a spurious connection between two neurons (Post_1_ and Post_2_) receiving common excitatory drive from a single presynaptic neuron (Pre) with different delays (orange cross-correlogram) B) Scatter plots show normalized spike “transmission” probabilities from different sets of simulations where the true connections to the postsynaptic neurons have different types of short-term synaptic plasticity (both Depressing, both Facilitating, and one Depressing; one Facilitating). Lines with same colors as scatter plots show the estimated spike probability from the TM-GLM. Here the data and model fits are averaged across 150 rounds of simulations (50 for each combination) and are normalized in order to have a spiking probability of 1 for the longest ISIs C) We then fit spike “transmission” probability for 38 pairs of neurons from the multi-electrode array (MEA) recording where there was no clear monosynaptic connection (putative non-connections). Observed (left) spike transmission probabilities show relatively little variation as a function of one neuron’s ISIs, but the TM-GLM (right) does describe what variation there is. Insets show example cross-correlograms from two of these putative non-connections.

We also fit the TM-GLM to several (n=38) pairs of neurons from the MEA data all with average firing rates in range of 3-15 Hz and where there was no clear peak in the cross-correlogram (0-5ms following the spikes of one neurons). In these cases, although the coupling filter is likely fitting noise and does not describe a realistic synaptic effect (median latency 0.7 ms, median time-constant 0.02 ms), the TM-GLM does describe small variations in the ISI-dependent pattern of spike “transmission” probability (Fig 8C). These patterns are not as pronounced as the patterns observed in the identified and putative monosynaptic connections described above, but they also appear to have structure that the TM-GLM can account for. Altogether, these results illustrate how the TM-GLM simply aims to account for short-term dynamics in the spiking probability of one neuron in reference to the spikes of another neuron. Correctly identifying monosynaptic connections is a necessary first step before the short-term dynamics can be meaningfully interpreted.

## Discussion

Here we developed a dynamic model of functional connectivity, the TM-GLM, and applied this model to disentangle synaptic and nonsynaptic contributions to excitatory spike transmission in vivo. Short-term synaptic plasticity (STP) has been extensively studied with intracellular recordings where the amplitudes of individual postsynaptic potential/currents (PSP/PSCs) can be directly measured. However, the relationship between STP and *in vivo* spike transmission patterns is complex. Patterns of postsynaptic spike transmission are highly diverse and multiple factors beyond STP shape these patterns (Swadlow and Gusev, 2001; English et al., 2017). Here, using a model-based approach, we characterized these diverse spike transmission patterns at identified and putative excitatory synapses and attribute this diversity to different combinations of short-term synaptic plasticity, synaptic summation, and post-spike history effects. We then showed how this modeling framework has the potential to capture stimulus-specific and cell-type-specific changes in spike transmission *in vivo*.

Estimating static functional connectivity using spike times has revealed network structure in the retina (Pillow et al., 2008) and hippocampus (Harris et al., 2003), can reconstruct true physiological circuitry (Gerhard et al., 2013), and improves encoding and decoding (Truccolo et al., 2005; Pillow et al., 2008; Stevenson et al., 2012). However, synaptic weights can change dramatically over time and can also depend on external stimuli and behavior (Fujisawa et al., 2008). Although, standard GLMs can partially capture the first-order effects of recent presynaptic spikes on postsynaptic spiking probability, they fail to capture the nonlinear dynamics of synaptic transmission affected by longer sequences of presynaptic spikes. With a static coupling term the GLM can account for the average change in the postsynaptic spiking probability following a presynaptic spike, but it does not make detailed predictions about the variations in this probability. Here we show that, by including a dynamical model of short-term plasticity, we can capture diverse pattern of spike transmission probability and substantially improve prediction of postsynaptic spiking. In a recording from the endbulb of Held (ANF-SBC) we further found that spike transmission patterns differed between stimuli, and that these differences were well-described by a single TM-GLM. Although the STP-parameters were the same for all stimuli, the different presynaptic spike patterns yield different patterns of spike transmission. Since spike transmission probability in the TM-GLM depends on the full history of presynaptic spiking, this model can account for changes on behavioral timescales even in the absence of adaptation or other forms of plasticity (e.g. STDP, LTP). Using the models for the short-term dynamics of spike transmission estimated in one setting we may also be able to more accurately predict responses to novel presynaptic patterns and, in sensory systems, novel stimuli.

Previous *in vitro* studies have shown that STP dynamics depend on both presynaptic and postsynaptic cell-types (Thomson and Lamy, 2007). Using a large multi-electrode recording from a freely behaving mouse, we investigated the dynamics of synaptic connections from putative excitatory neurons to two different subtypes of putative inhibitory neurons: putative fast-spiking (FS) and putative regular-spiking (RS). Using only spike times, we find that spike transmission shows slightly higher facilitation for excitatory-RS compared to the excitatory-FS connections. Although drawing strong conclusions about the parameters of the model is difficult due to potential confounds, the STP dynamics reflect this same pattern and are in line with previous *in vitro* findings (Thomson and Lamy, 2007). Including short-term dynamics into the model also significantly improves the prediction of postsynaptic spiking. As large-scale extracellular recording techniques advance, models such as the TM-GLM may allow us to characterize and compare the short-term dynamics of spike transmission of many different cell types, brain regions, and species.

Several details of the model may impact our results. Here we employed an extended GLM with a logistic spike nonlinearity, since it appears to better describe strong connections, such as the ANF-SBC, better than the traditional exponential nonlinearity. However, other nonlinearities may be better for other neurons (McFarland et al., 2013). There are also alternatives to the Tsodyks-Markram model for modeling synaptic dynamics (Hennig, 2013). Although the TM model is biologically plausible, it only tracks average, deterministic dynamics of postsynaptic potentials, while ignoring the stochasticity of synaptic release (Barri et al., 2016; Bird et al., 2016). Finally, there are many covariates that could be added to improve model performance, including local field potentials (Kelly et al., 2010), connections to other simultaneously observed presynaptic neurons (Harris et al., 2003), higher-order history or coupling terms (Robinson et al., 2016; Song et al., 2018), and covariates related to other types of plasticity (Stevenson et al., 2011; Linderman et al., 2014; Robinson et al., 2016; Amidi et al., 2018; Bayat Mokhtari et al., 2018). Despite these simplifying assumptions and the fact that we only observe a fraction of inputs to the neuron, the TM-GLM captures a wide diversity of *in vivo*, excitatory spike transmission patterns.

Although our model provides a tool to characterize the dynamics of spike transmission, there may be fundamental limitations to how well true synaptic dynamics can be estimated from spike observations. Firstly, functional connections inferred from spikes do not necessarily guarantee anatomical connections. A peak in the cross-correlogram does not conclusively indicate the presence of a monosynaptic connection (Moore et al., 1970). In most cases, we assume that the transient, short-latency increase in postsynaptic spiking activity following a presynaptic spike indicates the presence of an excitatory monosynaptic connection (Perkel et al., 1967). Nevertheless, verifying connections using optogenetics (English et al., 2017), juxtacellular recordings (Pinault, 2011), or imaging (Weiler et al., 2008) may provide more confidence in determining true monosynaptic connections. Secondly, we employ a spiking model that does not explicitly account for the detailed membrane potential of the postsynaptic neuron. Although there are links between the GLM and voltage-based models (Latimer et al., 2014, 2018), other approaches to modeling synaptic transmission with realistic spike-generation mechanisms, currents, and even dendritic morphology may more accurately reflect subthreshold dynamics (Ladenbauer et al., 2018). Thirdly, long-term changes in the synaptic weight may alter the short-term dynamics. Experiments show that short-term depression may be reduced after long-term depression and increased after long-term potentiation (Markram and Tsodyks, 1996; Sjöström et al., 2007; Costa et al., 2015, 2017). Accounting for these long-term changes in synaptic strength may allow for more accurately estimation of STP. Finally, there are many other factors that are likely to affect short-term spike transmission dynamics including, dendritic spikes (Bono and Clopath, 2017), receptors nonlinearities (Magee, 2000), such as those in NMDA receptors, changes in spike threshold due to sodium inactivation (Naud et al., 2011) or coupled to the subthreshold activity (Mensi et al., 2016), feed-forward inhibition (Pouille and Scanziani, 2001), feedback inhibition (Suzuki and Bekkers, 2012), or disinhibition (Letzkus et al., 2015). With intracellular observations these effects can generally be separated from the synaptic dynamics based on the timing of the signals. However, since these effects directly alter spike timing, they may act as confounders for models based on spike observations. Although they could potentially be incorporated in future models, omitting these effects from the model presented here may result in biased parameter estimates for both the synaptic and non-synaptic effects that are included (Stevenson, 2018).

Intracellular observations in controlled settings have found that short-term synaptic dynamics vary depending on the pre- and postsynaptic cell type (Thomson and Lamy, 2007; Lee et al., 2019) as well as brain region (Dittman et al., 2000; Wang et al., 2006), age (Reyes et al., 1998), and species (Testa-Silva et al., 2014). Additionally, short-term synaptic dynamics appear to vary with stimulus type and the larger computational function of the neural circuit (Karmarkar and Buonomano, 2007). To link synaptic dynamics to circuit-level neural computations we will need to study these dynamics during natural ongoing activity (Klyachko and Stevens, 2006) and ultimately during natural behavior. Since short-term synaptic plasticity affects not only the postsynaptic membrane potential but also the probability of postsynaptic spiking (Markram et al., 1998; Swadlow and Gusev, 2001; London et al., 2002; English et al., 2017), it may be possible to indirectly observe the effects of synaptic dynamics on spike transmission. Here we examined this possibility by including the effects of short-term synaptic plasticity in models of functional connectivity. Using this approach, we characterized diverse, stimulus-dependent, and cell-type-specific patterns of excitatory spike transmission using spike observations alone.

## Data and software availability

All data and software central to the conclusion of this study are available at https://github.com/abedghanbari2/TM-GLM.

## Acknowledgements

Thanks to Nick Steinmetz for providing MEA dataset. AG, NR, and IHS were supported by NSF CAREER 1651396 to IHS. BE was partially funded via an NWO Open Grant (ALWOP.346) and an NWO VIDI grant (016.Vidi.189.052).

